# A cross–nearest neighbor/Monte Carlo algorithm for single-molecule localization microscopy defines interactions between p53, Mdm2, and MEG3

**DOI:** 10.1101/857912

**Authors:** Nicholas C. Bauer, Anli Yang, Xin Wang, Yunli Zhou, Anne Klibanski, Roy J. Soberman

**Affiliations:** Division of Nephrology, Department of Medicine, Massachusetts General Hospital and Harvard Medical School, Charlestown, Massachusetts, United States; Neuroendocrine Unit, Department of Medicine, Massachusetts General Hospital and Harvard Medical School, Boston, Massachusetts, United States

**Author notes:** These authors contributed equally to the work (Anli Yang, Xin Wang). These authors contributed equally to the work (Anne Klibanski, Roy J. Soberman). Department of Breast Oncology, Sun Yat-sen University Cancer Center, Guangzhou, Guangdong, China (Anli Yang). Department of Histology and Embryology, Guangdong Pharmaceutical University, Guangzhou, Guangdong, China (Xin Wang). Corresponding author: Roy Soberman.

**Keywords:** p53, mouse double minute 2 homolog (Mdm2), long noncoding RNA (long ncRNA, lncRNA), microscopy, computational biology, image analysis, single-molecule localization microscopy (SMLM), stochastic optical reconstruction microscopy (STORM)

## Abstract

The functions of long noncoding (lnc)RNAs such as MEG3 are defined by their interactions with other RNAs and proteins. These interactions, in turn, are shaped by their subcellular localization and temporal context. Therefore, it is important to be able to analyze the relationships of lncRNAs while preserving cellular architecture. The ability of MEG3 to suppress cell proliferation led to its recognition as a tumor suppressor. MEG3 has been proposed to activate p53 by disrupting the interaction of p53 with Mdm2. To test this mechanism in the native cellular context, we employed two-color direct stochastic optical reconstruction microscopy (dSTORM), a single-molecule localization microscopy (SMLM) technique to detect and quantify the localizations of p53, Mdm2, and MEG3 in U2OS cells. We developed a new cross-nearest neighbor/Monte Carlo algorithm to quantify the association of these molecules. Proof of concept for our method was obtained by examining the association between FKBP1A and mTOR, MEG3 and p53, and Mdm2 and p53. In contrast to previous models, our data support a model in which MEG3 modulates p53 independently of the interaction with Mdm2.

## Introduction

Long noncoding RNAs (lncRNAs) function in cell-type and subcellular localization-dependent contexts; how they do so is incompletely understood. The human *MEG3* lncRNA gene is located on chromosome 14q32 and belongs to the conserved, imprinted *DLK1-MEG3* locus (1,2). MEG3 transcripts are detected in a wide range of normal tissues, including endocrine tissues, brain, gonads, and placenta (1). MEG3 modulates the activity of multiple miRNAs; for example, MEG3 functions as a decoy for miR-138 (3) allowing it to regulate the generation of IL-1β in macrophages in models of host defense. MEG3 has also been reported to directly interact with DNA to modulate the transcription of TGF-β pathway genes (4).

Based on the observation that *MEG3* expression is lost in clinically non-functioning pituitary adenomas, we identified *MEG3* as a tumor suppressor (1,5–7). Compared to normal tissue, MEG3 expression is also significantly reduced or absent in meningiomas (8), epithelial ovarian cancer (9), and squamous cell carcinoma of the tongue (10); supporting its role as a tumor suppressor. This function was further supported by studies of tumor xenograft growth in nude mice (11,12). Several studies demonstrated that MEG3 expression causes an increase in cellular tumor antigen p53 (p53, UniProtKB P04637) levels and selectively activates p53 target gene expression (11,13–16), suggesting that MEG3 exerts its cellular functions via p53. How the MEG3 lncRNA activates p53 remains unsolved.

p53 coordinates a transcription program to stall the cell cycle, promote DNA repair, and initiate senescence or apoptosis (17). The primary modulators of p53 activity are the E3 ubiquitin-protein ligase Mdm2 (Mdm2, UniProtKB Q00987) and its heterodimer partner protein Mdm4 (Mdm4, UniProtKB O1515) which constitutively polyubiquitinate p53 for proteasomal degradation, maintaining p53 at low levels (18–20). Thus, modulating the p53–Mdm2/4 interaction is a critical point of regulation for p53 activity. Signal-dependent post-translational modification of p53, including phosphorylation and acetylation, can block Mdm2/4 from binding to p53 and prevent its degradation (21). Stabilization of p53 may also be achieved through interaction with other proteins such as peptidyl-prolyl cis-trans isomerase NIMA-interacting 1 (Pin1) (22,23). It has been shown that MEG3 and p53 can be pulled down in one complex by immunoprecipitation (24,25). Therefore, one possible mechanism for p53 activation by MEG3 is to disrupt the p53–Mdm2/4 interaction.

Identifying molecular associations within the spatial context of the cell is necessary to fully define the behavior of MEG3. Single-molecule localization microscopy (SMLM) is exceptionally well positioned to provide this information. SMLM is unique from other microscopy approaches in that it provides high-accuracy coordinates of the positions of fluorophores rather than an image (although an image may be reconstructed from these localizations). As such, SMLM data must be analyzed with very different methods from traditional microscopy data which are still under active development (26). The first techniques applied traditional fluorescence image analysis approaches to the reconstructed images, although much of the unique information obtained by SMLM is lost this way. More promising approaches have looked at cluster-based and tessellation-based analyses (27,28), enabling the examination of supramolecular assemblies. However, there has been little work done towards using SMLM data to measure single molecular interactions.

To fully understand the interactions of MEG3 with p53 and to test the hypothesis that MEG3 disrupts p53–Mdm2 binding, we developed a new cross-nearest neighbor/Monte Carlo algorithm to quantify the association between molecules from direct stochastic optical reconstruction microscopy (dSTORM) data. We characterized the behavior of this method *in silico* and demonstrated a proof of concept by examining the association between FKBP1A and mTOR, MEG3 and p53, and Mdm2 and p53. In contrast to previous models, our data support a model in which MEG3 modulates p53 independently of Mdm2. Future work will build on this technique to examine the relationships of MEG3 to other cell components.

## Results

### Quantifying macromolecular associations by SMLM

We developed an SMLM approach that allowed us to identify potentially interacting macromolecules by calculating the probability that two localizations were anomalously close. We applied a Monte Carlo estimation method (Figure 1) that accounts for the local density around a candidate binding pair. This method is partly based on a technique recently introduced for examining the association of sparse mRNAs in neurons (29) but with substantial adaptations to deal with the unique properties of SMLM data.

**Figure 1.**
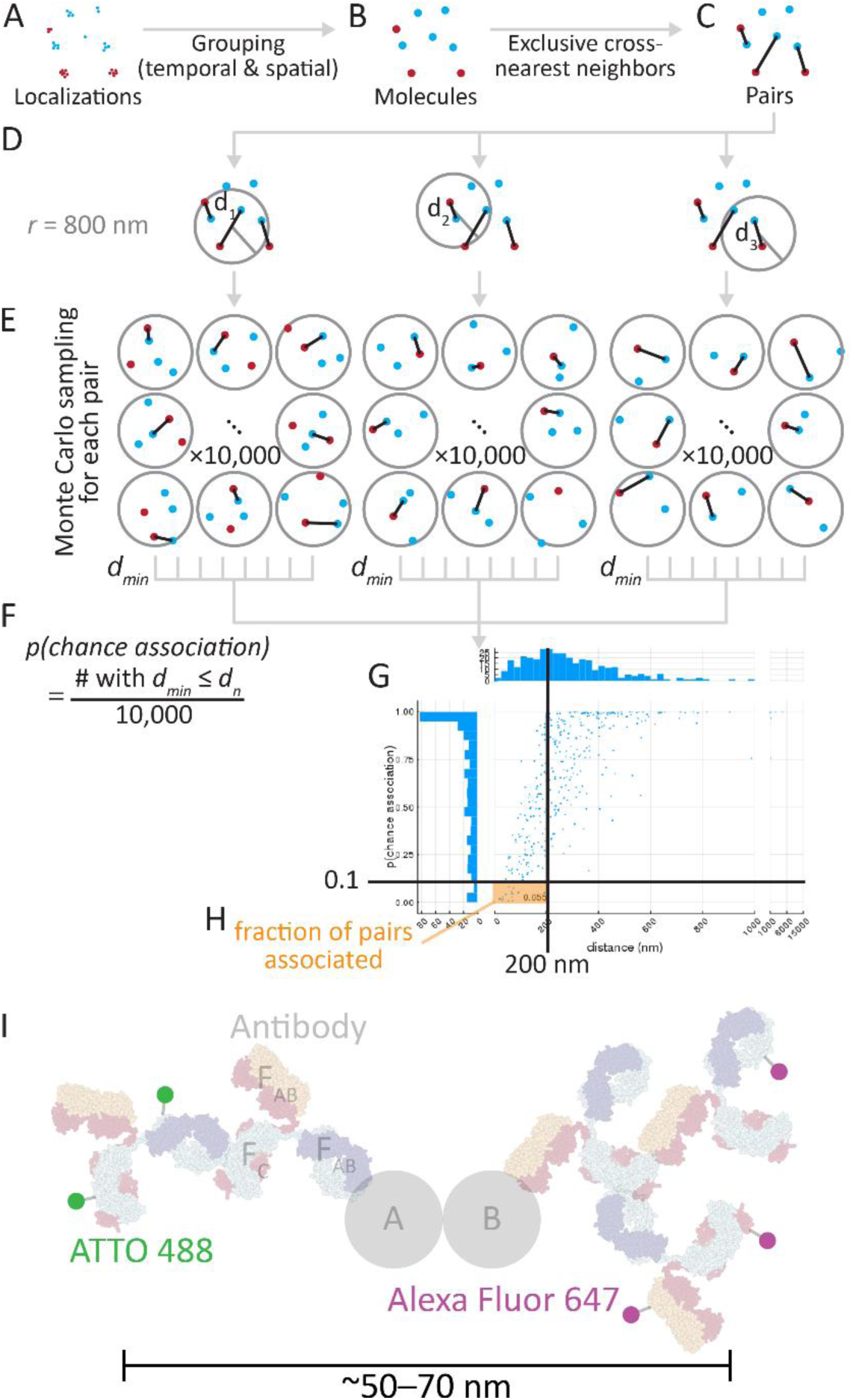
Cross-nearest neighbor/Monte Carlo method to estimate the fraction of molecules associated. Scattered localizations (A) were grouped over time and space to into “molecules” which have a position that is the average of their component localizations (B; see Figure S5). These molecules were exclusively paired to their cross-nearest neighbors (C). For each pair (D), 10,000 permutations of the molecules within radius *r* (800 nm) of the centroid of the pair were generated and the closest intermolecular distance was measured (E). The fraction of events less than the pair’s distance was the probability of chance association *(p(chance association))* (F). These values were accumulated across the whole cell (plotted in G), and the fraction of pairs with a probability of chance association < 0.1 and within a physically possible binding distance (< 200 nm), the fraction associated, was calculated (H). (I) The physical arrangement of a bound pair and antibody stack. Typical immunofluorescence uses expensive, target-specific primary antibodies and cheap secondary antibodies conjugated to a fluorophore like ATTO 488 and Alexa Fluor 647. For orientation, one antibody is labeled for its constant domain (F_C_) and two antigen binding domains (F_AB_). The physical size and arrangement mean that ~50-70 nm may separate signal from the two fluorophores when detecting a binding interaction between proteins A and B, and multiple fluorophores can produce signal spread over tens of nanometers. Antibody graphic was created using NGL Viewer (56) from RCSB PDB 1IGT.

In the first step nearby localizations were grouped into “molecules” using spatial and temporal thresholds (Figure 1A and 1B). A characteristic of dSTORM is that there is no guarantee that a single molecule will be represented by a single localization. Consider the p53 tetramer: it may be bound by up to 4 primary single-epitope antibodies, each of which may be bound by 1 or 2 secondary antibodies, each of which may have 0 to 8 fluorophores attached (despite the average being ~1 dye molecule/antibody), and each fluorophore may blink many times before permanent bleaching. A grouping algorithm is important for dSTORM data to remove such autocorrelated localizations for our downstream analysis, which here assumes that each molecule’s location is independent of each other molecule.

Second, “molecules” from each channel were paired together through an exclusive cross-nearest neighbor algorithm: closest pair identified then removed, repeating until all possible pairs were made (Figure 1B and 1C). The resulting list of pairs is guaranteed to contain all detectable binding events.

The third phase of the algorithm assesses the probability that each pair is associated by chance. Within the local neighborhood (radius *r* = 800 nm, Figure 1D), 10,000 random permutations of the positions of the molecules within this radius were generated and the smallest paired distance measured in each iteration (Figure 1E). The fraction of permutations in which a distance *d_n_* less than or equal to the observed distance *d_min_* was recorded as the probability of chance association (*p(chance association)*) for that molecule pair (Figure 1F). These steps were repeated for each pair in each cell, and a graph of distance and probability of chance association may be generated (Figure 1G). This plot from a representative cell shows that larger distance is correlated with higher probability of chance association, with wide variability due to local density changes.

Finally, these pairwise measures of association were reduced into a summary value which would correlate with fraction bound. We considered pairs with a probability of chance association less than 0.1 and a distance of less than 200 nm to be associated, and used that value to generate the final output, fraction associated (Figure 1H). In this example, the average distance of the pairs in the associated fraction is approximately 50 nm, which corresponds well with the range expected due to the size of the antibody stacks used to detect molecules (up to ~50-70 nm between fluorophores, Figure 1I). The largest expected distances would come from the p53 tetramer, which would potentially add another 5-10 nm to the distance between fluorophores but well within the 200 nm cutoff chosen. In this dataset, pairs with a large distance but a low probability of chance association were rare; most of those pairs classified as unassociated were due to moderately close pairs in dense areas.

### Algorithm performance improves with lower density, shorter distances

A natural limitation of this algorithm is that it strongly depends on the density of localizations and the distance between the fluorophores of an associated pair. To characterize this behavior, we simulated 30 distributions of molecules (i.e. the product of the grouping algorithm described above) for each combination of density (100, 200, 500, 1000, 2000, 5000, and 10000 molecules of each kind within a 250 μm^2^ circle), percent binding (0, 1, 2, 5, 10, 20, 50, and 100%), and physical distance between fluorophores (10, 20, 50, 100, and 200 nm). Higher density naturally means that the two sets of localizations are closer together on average (Figure 2, columns from left to right), while larger binding distance inflates the value they converge to as more of the molecules are bound together (Figure 2, rows from top to bottom). The Monte Carlo component of the association algorithm adjusts for local density around a putative associated pair, but the algorithm’s sensitivity is reduced by higher global density (Figure 3, columns from left to right). Larger physical distance between fluorophores in a bound pair similarly reduces sensitivity, as the pair becomes harder to distinguish from the background distribution (Figure 3, rows from top to bottom). Normalizing the fraction associated using 0% and 100% mean values improves the correspondence between true fraction bound and measured fraction associated, but the measured value becomes a severe underestimate of the true fraction bound at higher densities and increased pair distances and can produce hard to interpret negative values (Figures S1 and S2). We chose not to apply this simulated normalization approach to the biological data, though a similar approach could be useful in an advanced iteration of our method.

**Figure 2.**
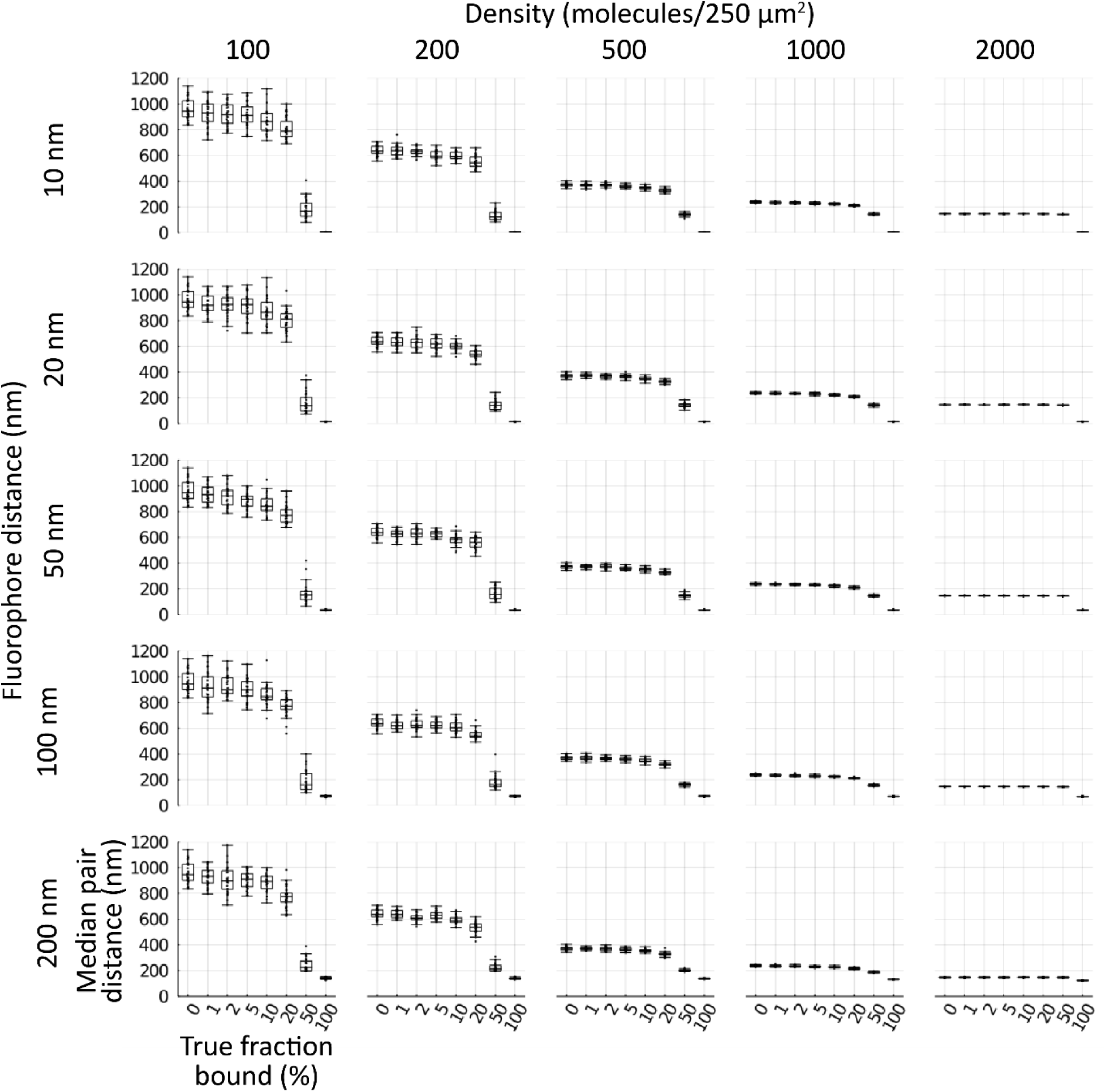
Density and pair distance determine median distance. Simulated cells of 100, 200, 500, 1000, 2000 molecules of each type (columns left to right) within a 250 μm^2^ circle with 10, 20, 50, 100, 200 nm separation between pairs (rows top to bottom) for cells with 0, 1, 2, 5, 10, 20, 50, or 100% binding were generated (n = 30 each condition). Molecules for each simulated cell was run through the cross–nearest neighbor/Monte Carlo algorithm and median distance between pairs measured. Boxes indicate median +/- upper and lower quartile; whiskers indicate the range excluding outliers.

**Figure 3.**
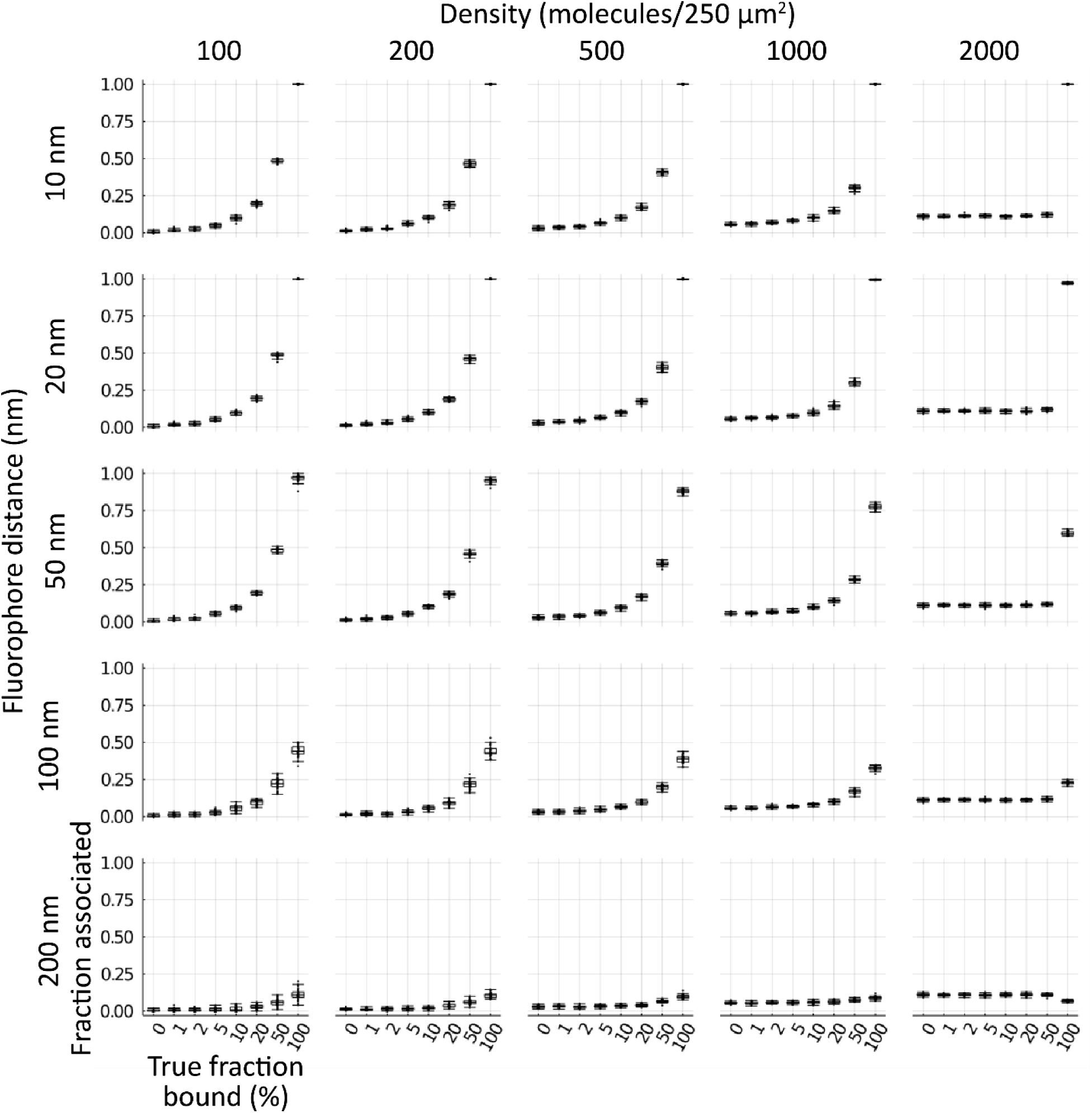
Density and pair distance influence fraction associated. Simulated cells of 100, 200, 500, 1000, 2000 molecules of each type (columns left to right) within a 250 μm^2^ circle with 10, 20, 50, 100, 200 nm separation between pairs (rows top to bottom) for cells with 0, 1, 2, 5, 10, 20, 50, or 100% binding were generated (n = 30 each condition). Molecules for each simulated cell were run through the cross–nearest neighbor/Monte Carlo algorithm and median distance between pairs measured. Boxes indicate median +/- upper and lower quartile; whiskers indicate the range excluding outliers.

### Detection of FKBP1A-mTOR interaction

In order to characterize the algorithm in a known biological system, we chose to look at the well-characterized rapamycin-induced binding of Peptidyl-prolyl cis-trans isomerase FKBP1A (FKBP1A, UniProtKB P62942) and Serine /threonine kinase mTOR (mTOR, UniProtKB P42345) (30). U2OS osteosarcoma cells were treated with 0 or 10 ng/mL rapamycin for 24 hours then fixed. FKBP1A was labeled with a secondary antibody conjugated with Alexa Fluor 647 (magenta) and mTOR was labeled with a secondary antibody conjugated to ATTO 488 (green). Large tiled widefield fluorescence images were taken (Figure S3) and ten individual cells were randomly selected from these fields for dSTORM analysis, in each of five replicates (Figure S4). Representative cells are shown in Figure 4A, widefield (left three columns) and dSTORM localizations and grouped molecules (right two columns, respectively). Expected strong nuclear mTOR fluorescence and cytoplasmic FKBP1A fluorescence is apparent, with no large-scale change upon rapamycin treatment. As seen in the detail of the molecule groups (right column), the grouping operation is slightly biased towards merging nearby clusters.

**Figure 4.**
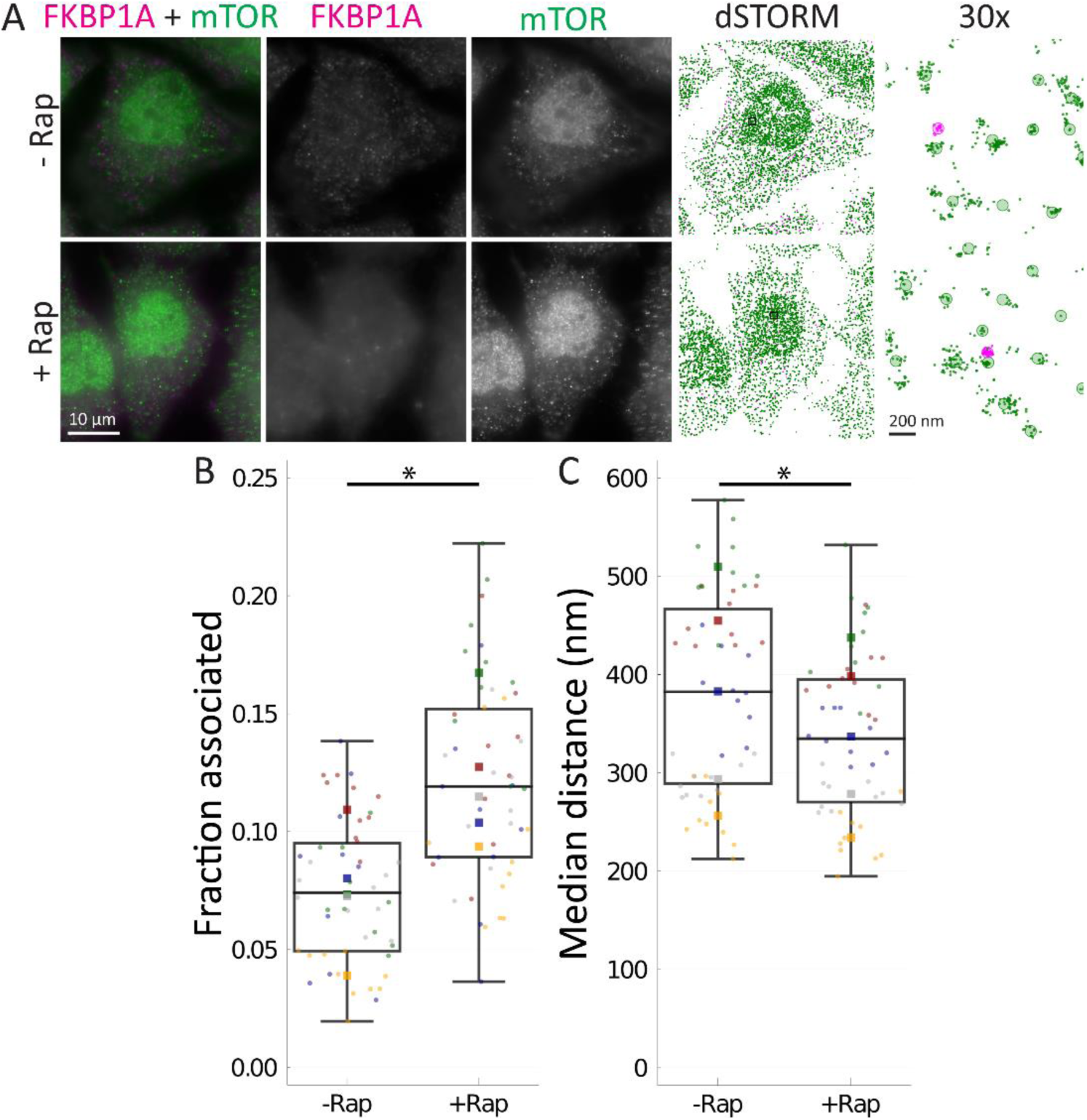
Method detects rapamycin-induced FKBP1A association with mTOR. U2OS-MEG3 cells were treated for 24 h with or without 10 ng/mL rapamycin. (A) Cells were stained for mTOR with a secondary antibody conjugated to ATTO 488 (green) and for FKBP1A with a secondary antibody conjugated to Alexa Fluor 647 (magenta). From left to right: Merged image; RNA channel; p53 channel; dSTORM localization map; 30x inset of dSTORM localizations in the black box, with shaded circles indicating “molecules”. Scale bars are 10 μm, or 200 nm (right column). (B) Fraction of pairs associated, as defined by a probability of chance association < 0.1 (i.e., correction for local density) and distance < 200 nm (upper limit for binding distance, accounting for error). (C) Median distance between pairs for each cell (nm). Boxes indicate median +/- upper and lower quartile; whiskers indicate the range excluding outliers. Data points are colored by replicate. For each condition, single molecule localizations were collected from 10 randomly chosen cells in 5 separate experiments. Means for each replicate are indicated by same-colored squares. * indicates p < 0.05 by ANOVA.

Using our cross-nearest neighbor/Monte Carlo method, we were able to detect a strong, significant increase in fraction associated of FKBP1A-mTOR pairs with rapamycin treatment, from 0.0749 ± 0.02504 to 0.121 ± 0.0286, by ANOVA with rapamycin treatment within replicates (F = 11.83, p = 0.02630, ω^2^ = 0.2687) (Figure 4). The 0 ng/mL data make clear that the method does not measure direct binding, but association; factors other than binding interactions can lead to associations between molecules. Instead, we suggest the data produced by this method be interpreted as measuring relative differences in association, and that data from a single condition is probably insufficient independent evidence to determine if two molecules are associated. The strength of rapamycin-induced binding leads to the question of why the fraction association with rapamycin present is not higher. The fraction of pairs of detected molecules may not represent the total number of pairs available for binding. Pair members may be held apart by compartmentalization, or binding may be blocked by other factors. With this interaction, mTOR is present in two separate complexes, mTOR complex 1 and 2 (MTORC1/2); MTORC1, but not MTORC2, may be bound by FKBP1A due to steric hindrance caused by other components in MTORC2 (30). Also as noted in the previous section, dense labeling and lengthy distances between fluorophores may also lead to under-detection of true associations.

For comparison, we also applied a naïve median distance approach, where we calculated the median of the pairwise distances for each cell (Figure 4). In this simple approach, there was a high degree of overlap between conditions and a significant but moderate decrease in median distance, from 380 ± 106.3 nm to 337 ± 83.4 nm (F = 15.97, p = 0.01618, ω^2^ = 0.04774) (Figure 4). Thus, our cross-nearest neighbor/Monte Carlo-based approach provides a stronger and more robust measure of association over simpler approaches.

### MEG3 associates with p53 and is insensitive to MEG3 induction

We developed U2OS osteosarcoma cell clones containing a doxycycline-inducible MEG3 (U2OS-MEG3) and confirmed that MEG3 was induced 100-200-fold on doxycycline treatment as determined by qRT-PCR. We established conditions for performing dSTORM, simultaneously imaging RNA using fluorescence *in situ* hybridization (FISH) and proteins by immunofluorescence (IF). After 20 h induction with doxycycline, cells were fixed and MEG3 was labeled with a tiled probe set conjugated with Quasar 670 (magenta), and p53 was labeled with a secondary antibody conjugated to ATTO 488 (green). Cells were separately labeled for *GAPDH* mRNA with a tiled probe set conjugated with Quasar 570 (magenta) as a negative control. Large tiled widefield fluorescence images were taken (Figure S5) and ten individual cells were randomly selected from these fields for dSTORM, in each of three replicates (Figure S6). Representative cells are shown in Figure 5, widefield (left three columns) and dSTORM localizations and grouped molecules (right two columns, respectively). The intense fluorescence indicating MEG3 is readily apparent in the nucleus of the cells treated with doxycycline, along with p53, while very little MEG3 is apparent in untreated cells (Figure 5A). Doxycycline treatment had no apparent effect on the GAPDH mRNA distribution (Figure 5B).

**Figure 5.**
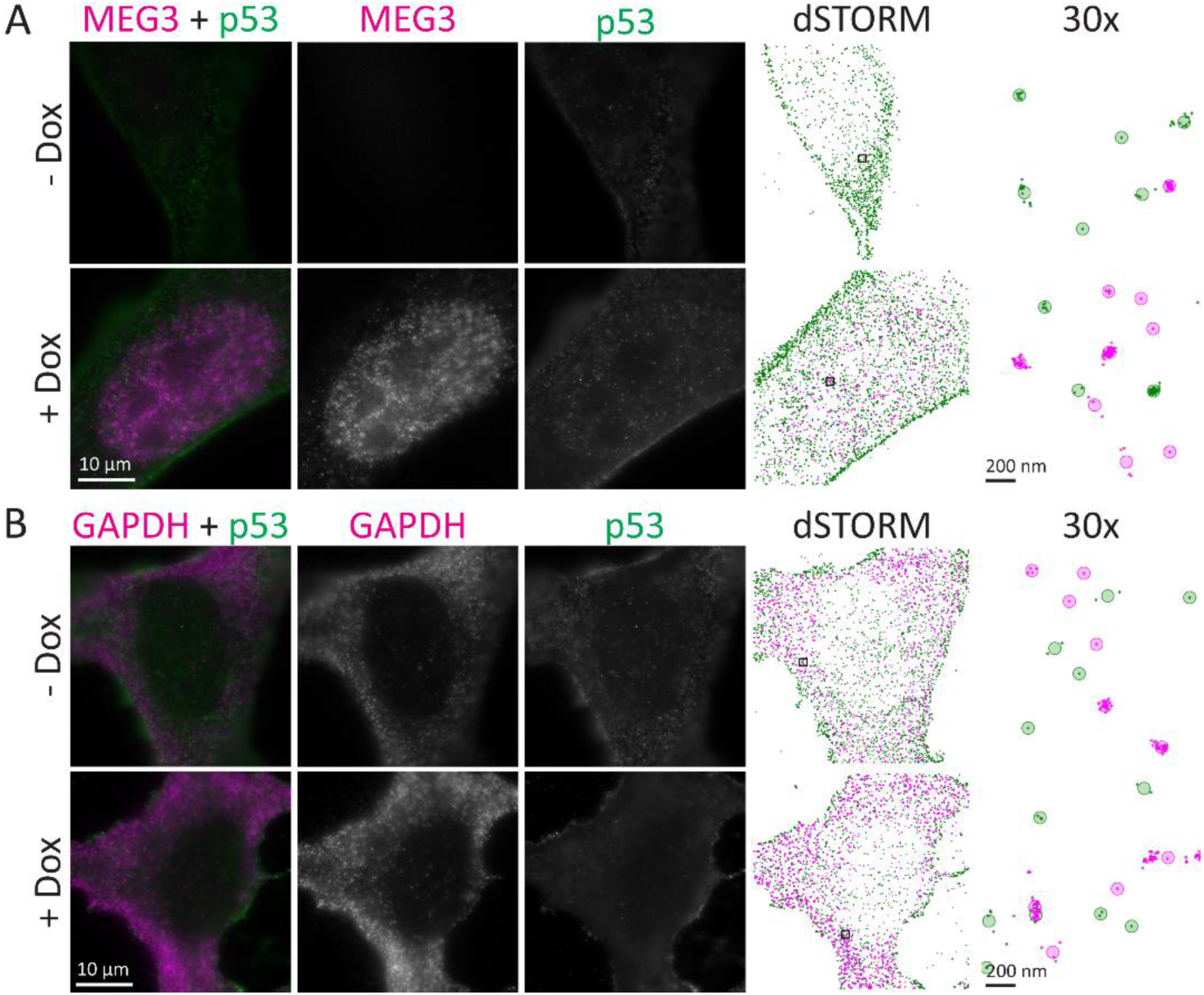
MEG3 localized in the nucleus of U2OS cells by epifluorescence and 2-color dSTORM. *MEG3* was induced by treatment of U2OS-MEG3 cells for 20 h with 1 μg/mL doxycycline. Cells were stained for p53 with a secondary antibody conjugated to ATTO 488 (green) and either tiled oligonucleotides recognizing MEG3 (Quasar 670, magenta, A) or GAPDH (Quasar 570, magenta, B). From left to right: Merged image; RNA channel; p53 channel; dSTORM localization map; 30x inset of dSTORM localizations in the black box, with shaded circles indicating “molecules”. Scale bars are 10 μm, or 200 nm (right column).

Using our cross-nearest neighbor/Monte Carlo method, we found a stark difference between MEG3 and *GAPDH* mRNA in terms of fraction associated (Figure 6A). ANOVA of doxycycline treatment within replicates and between RNAs demonstrated that the fraction associated was significantly larger for MEG3 than for *GAPDH* mRNA (F = 14.41, p = 0.01917, ω^2^ = 0.3497). Since only the RNA main effect was significant, a followup one-way ANOVA was performed within each RNA type. For *GAPDH* mRNA, no change due to doxycycline was observed (-Dox (white box): 0.0430 ± 0.00312 vs. +Dox (gray box): 0.0422 ± 0.00770; F = 0.04628, p = 0.8496, ω^2^ = −0.03593) (Figure 6A, right). There are several reasons why the method may detect a very low level of association between p53 and *GAPDH* mRNA. Some level of background associations may be expected due to density of localizations. p53 is also known to have promiscuous non-specific RNA binding capacity (31). Another contributor may be crosstalk due to the spectral overlap of the fluorophore used for *GAPDH* mRNA and p53 (Quasar 570 vs. ATTO 488). However, we expect true molecular associations to lead to a much higher measured fraction associated.

**Figure 6.**
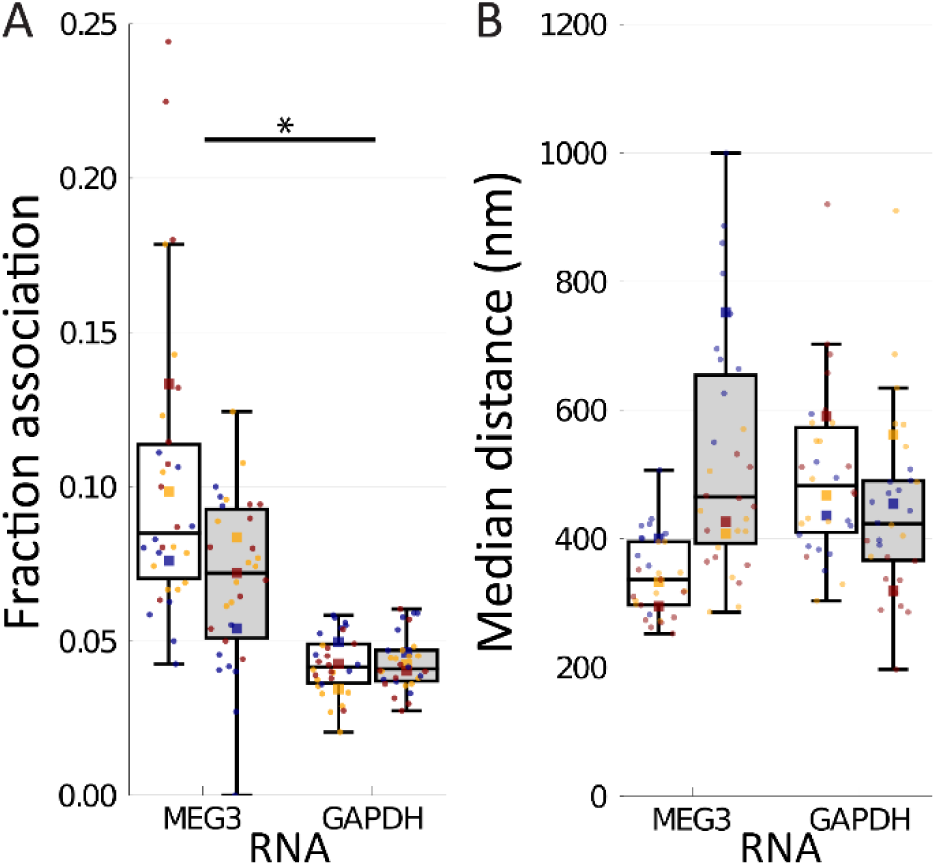
MEG3 associated with p53. *MEG3* was induced by treatment of U2OS-MEG3 cells for 20 h with (gray) or without (white) 1 μg/mL doxycycline. The cells were then fixed and stained for 2-color dSTORM of MEG3 and p53 (left) or *GAPDH* mRNA and p53 (right). For each condition, single molecule localizations were collected from 10 randomly chosen cells in 3 separate experiments. (A) Fraction of pairs associated, as defined by a probability of chance association < 0.1 (i.e., correction for local density) and distance < 200 nm (upper limit for binding distance, accounting for error). (B) Median distance between pairs for each cell (nm). Boxes indicate median +/- upper and lower quartile; whiskers indicate the range excluding outliers. Data points are colored by replicate. Means for each replicate are indicated by same-colored squares. * indicates p < 0.05 by ANOVA.

In contrast with *GAPDH* mRNA, the fraction of MEG3 associated with p53 was substantially higher than with *GAPDH* mRNA and doxycycline induction had a medium-large effect, but was not significantly different (-Dox (white box): 0.103 ± 0.0149, +Dox (gray box): 0.0700 ± 0.02888; F = 5.136, p = 0.1516, ω^2^ = 0.1212) (Figure 6A, left). A reduction or no change in fraction of MEG3 associated with p53 following MEG3 induction suggests that MEG3 binding to p53 stays roughly constant as the level of MEG3 increases. Since the fraction is based on the number of pairs and MEG3 is the limiting factor, little change in fraction bound when MEG3 expression is induced implies a substantial increase in the absolute number of associations between MEG3 and p53 in the cell. MEG3 also interacts with numerous other macromolecules, including Polycomb repressive complex 2 (PRC2) (32,33), which may absorb much of the increased levels of MEG3.

For comparison, we also applied a naïve median distance approach, where we calculated the median of the pairwise distances for each cell (Figure 6B). In this simple approach, there was a high degree of overlap between MEG3 and *GAPDH* mRNA distributions and no overall effect of RNA on median distance was found (487 ± 151.8 nm to 420 ± 105.0 nm; F = 0.2344, p = 0.6535, ω^2^ = −0.05747) (Figure 6B). These data support our cross-nearest neighbor/Monte Carlo-based approach as providing a more consistent and robust measure of association over simpler approaches.

### Mdm2-p53 binding maintains stable equilibrium

To determine whether MEG3 causes accumulation of p53 by disrupting the Mdm2-p53 interaction, MEG3 was induced by doxycycline in U2OS-MEG3 cells, with or without the addition of the Mdm2-p53 binding inhibitor nutlin-3a. After 24 h of treatment, the cells were fixed. p53 was labeled with a secondary antibody conjugated to Alexa Fluor 647 (magenta) and Mdm2 was labeled with a secondary antibody conjugated to ATTO 488 (green). Large tiled widefield fluorescence images were taken (Figure S7) and ten individual cells were randomly selected from these fields for dSTORM analysis, in each of three replicates (Figure S8). Representative cells are shown in Figure 7, widefield (left three columns) and dSTORM localizations and grouped molecules (right two columns, respectively). Intense nuclear p53 fluorescence is observed on treatment with nutlin-3a, without much apparent change due to doxycycline. Mdm2 levels and localization change little between conditions.

**Figure 7.**
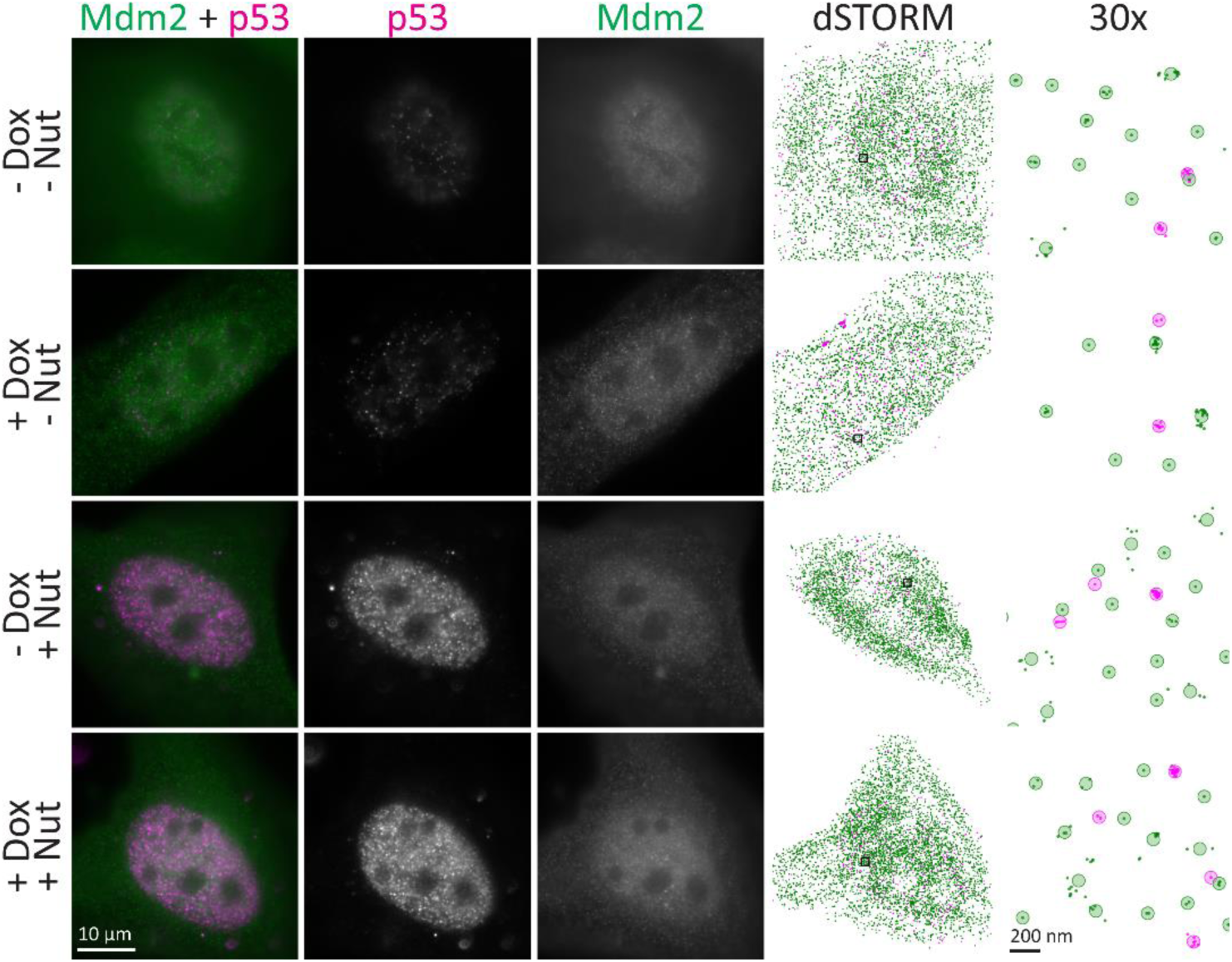
Nutlin-3a caused accumulation of p53 in the nucleus. *MEG3* was induced by treatment of U2OS-MEG3 cells for 20 h with 1 μg/mL doxycycline and/or 10 μM nutlin-3a. Cells were stained for Mdm2 with a secondary antibody conjugated to ATTO 488 (green) and for p53 with a secondary antibody conjugated to Alexa Fluor 647 (magenta). From left to right: Merged image; RNA channel; p53 channel; dSTORM localization map; 30x inset of dSTORM localizations in the black box, with shaded circles indicating “molecules”. Scale bars are 10 μm, or 200 nm (right column).

Using our cross-nearest neighbor/Monte Carlo method, we found that nutlin-3a treatment caused a large but not significantly different change in p53– Mdm2 association by ANOVA with nutlin-3a and doxycycline within replicates (-Nut: 0.0727 ± 0.01209, +Nut: 0.0517 ± 0.01703; F = 10.76, p = 0.08174, ω^2^ = 0.1289) (Figure 8A). Doxycycline treatment (MEG3 induction) had a small but not significantly different change (-Dox: 0.0565 ± 0.01230, +Dox: 0.0680 ± 0.02169; F = 3.265, p = 0.2125, ω^2^ = 0.02983). There was no significant interaction effect (F = 0.1912, p = 0.7046, ω^2^ = −0.05837). Importantly, these data suggest that MEG3 does not disrupt the overall level of p53– Mdm2 binding. However, nutlin-3a also did not definitively disrupt p53–Mdm2 binding by this measure. These observations may in part be explained through equilibrium. For example, nutlin-3a inserts into the pocket of Mdm2 that binds to p53, preventing Mdm2 from ubiquitinating p53 and marking it for proteasomal destruction (34). Thus, levels of both proteins may accumulate until they reach a high enough concentration to overcome the effect of nutlin-3a. It is also possible that the effects of nutlin-3a are more nuanced. Nutlin-3a has been shown to block Mdm2-mediated polyubiquitination and transrepression of p53 (35,36) and nutlin-3a causes Mdm2 to rapidly deplete from an artificial concentration of p53 (37) but does not block monoubiquitination and complexes were still observed by proximity ligation assay (36). These studies suggest that nutlin-3a may reduce high-affinity binding but not strongly affect transient interactions. Our results are consistent with this model.

**Figure 8.**
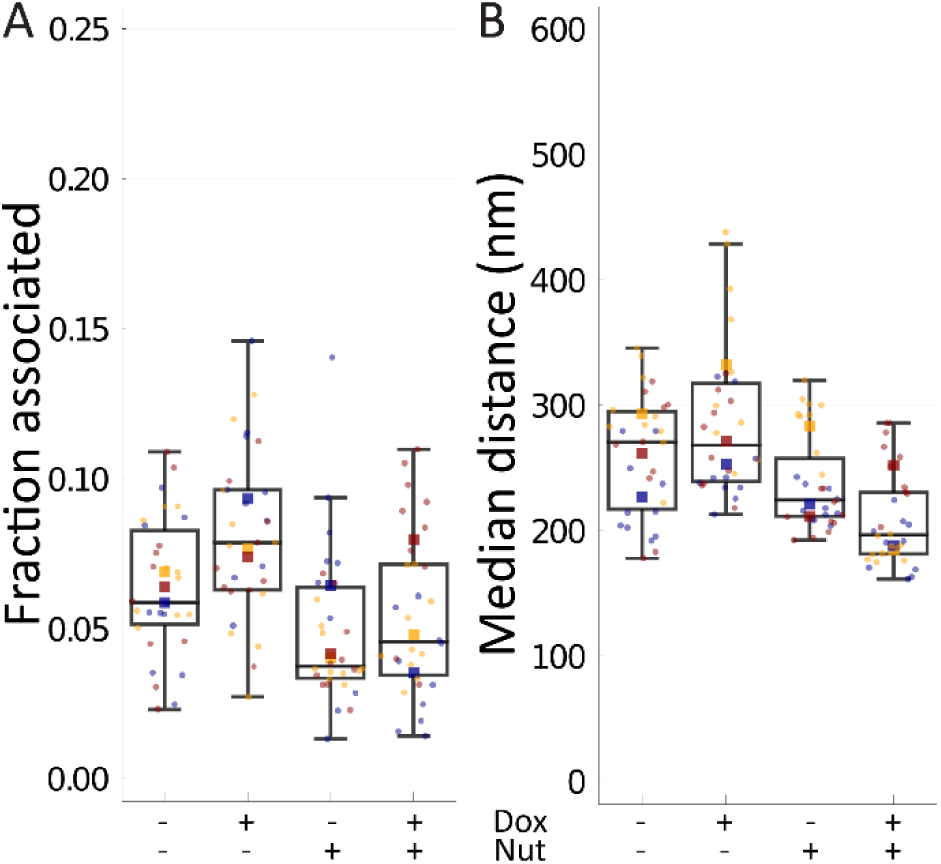
Neither nutlin-3a nor MEG3 significantly affected Mdm2-p53 binding. *MEG3* was induced by treatment of U2OS-MEG3 cells for 20 h with or without 1 μg/mL doxycycline and/or 10 μM nutlin-3a. Cells were then fixed and stained for 2-color dSTORM of p53 and Mdm2. For each condition, single molecule localizations were collected from 10 randomly chosen cells in 3 separate experiments. (A) Fraction of pairs associated, as defined by a probability of chance association < 0.1 (i.e., correction for local density) and distance < 200 nm (upper limit for binding distance, accounting for error). (B) Median distance between pairs for each cell (nm). Boxes indicate median +/- upper and lower quartile; whiskers indicate the range excluding outliers. Data points are colored by replicate. Means for each replicate are indicated by same-colored squares. * indicates p < 0.05 by ANOVA.

For comparison, we again conducted an analysis based on naïve median distances. As with MEG3-p53 binding, we found this simple approach produced overlapping distributions and less interpretable results. Nutlin-3a was associated with a large but not significant decrease in median distance (-Nut: 273 ± 36.2 nm, +Nut: 224 ± 38.2 nm; F = 11.60, p = 0.07643, ω^2^ = 0.2049) (Figure 8B). Doxycycline was not associated with any change in median distance (-Dox: 250 ± 34.6 nm, +Dox: 247 ± 55.2 nm; F = 0.02689, p = 0.8848, ω^2^ = −0.02940), and there was a small but nonsignificant interaction effect (F = 1.271, p = 0.3767, ω^2^ = 0.01809). The decrease in distance due to nutlin-3a is likely due to the increase in p53 expression levels when Mdm2 is inhibited from marking p53 for degradation.

## Discussion

We developed a mathematical approach to analyzing SMLM data that we used to interrogate the interactions of MEG3. Our overall approach takes advantage of high-resolution molecule position data to calculate distances between putative binding partners, assesses the probability that the two molecules are not associated, and thus provides an overall measure of the fraction of pairs of molecules likely associated. Using this technique, we distinguished between non-binding pairs *(GAPDH* mRNA-p53) and binding pairs (MEG3-p53, FKBP12-mTOR) inside cells and quantified the extent of association between Mdm2 and p53. The mechanism of MEG3 action suggested by our experiments is different than the previously proposed mechanism in which MEG3 acts by protecting p53 from polyubiquitination by Mdm2. Moreover, the fraction of association assessed between MEG3 and p53 indicates that there are insufficient stable interactions occurring to directly compete against p53–Mdm2 binding. These data suggest that MEG3 activates p53 through alternative mechanisms.

Under MEG3 induction, p53 transcription activation is selective, inducing certain p53 targets (e.g., *GDF15)* but failing to induce other p53 targets (e.g., *CDKN1A)* (16). A MEG3-p53 complex may not be competent to induce Mdm2 expression, thereby suppressing the negative feedback regulatory loop. MEG3 also interacts with the chromatin remodeler polychrome repressive complex 2 (PRC2) (32,33), which is responsible for forming heterochromatin at target sites. MEG3 targets PRC2 to certain sites via DNA triplex formation (e.g., TGF-β pathway genes) (4) and protects other sites from PRC2 activity (e.g., *MEG3* locus) (38). A recent investigation of MEG3 structure identified a pseudoknot in MEG3 critical for MEG3-dependent p53 activation, which however was not directly involved in p53 binding (39). It is also possible that MEG3 may modulate the activity or binding affinity of Mdm2 on p53 by forming a ternary complex with them. Similar interactions have been observed with p14^ARF^ (tumor suppressor ARF) (40), UCH-L1 (ubiquitin carboxyl-terminal hydrolase isozyme L1) (41), and the 5S RNP (42). Future work will need to address these alternative mechanisms. This algorithm could be expanded to consider ternary interactions to investigate these pathways.

There are some important challenges to the cellular SMLM-based association analysis we have developed, and which affect SMLM analysis approaches in general. First, despite the 10–20 nm resolution of each localization, the large distance between the molecule of interest and the fluorophore limit the analytical resolution. A typical two-antibody stack can have a displacement of up to ~35 nm from the bound epitope to the conjugated fluorophore; thus, the fluorophores for a bound pair of molecules may be separated by ~50-70 nm or more (Figure 1I), depending on the distance between epitopes. In addition, the antibody stack may be free to rotate and flex at the neck, adding variability in the position of the fluorophore during imaging (43). High density also impacts ability to differentiate low fraction associated, as illustrated in Figure 3. These distance issues may be addressed in part using nanobodies as secondary antibodies, which would reduce the distance to ~30–50 nm (Figure 9A). Nanobodies are small single-domain antibody fragments derived from camelids that are emerging as powerful and versatile tools for biology (44–46). Further distance reduction to ~10-30 nm and stable positioning could be achieved by introducing a fusion tag into the target gene and directly binding it with a nanobody (Figure 9B) (44). Further improvements to the algorithm may also be able to address these issues.

**Figure 9.**
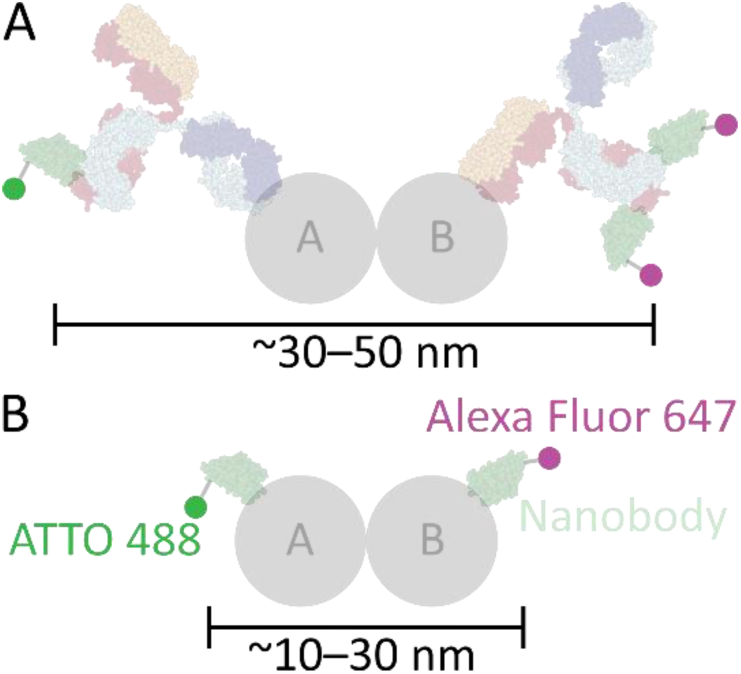
Nanobodies reduce distance between fluorophores for bound pairs. (A) Nanobodies (pale green) can be used in the place of secondary antibodies, reducing the distance to ~30–50 nm. (B) Nanobodies may also be used to bind directly to a target of interest, either native or via a small peptide fusion tag, reducing the distance to ~10–30 nm. Antibody and nanobody graphics were created using NGL Viewer (56) from RCSB PDB 1IGT and 5IVO, respectively.

A second set of limitations comes from the stochastic nature of SMLM. Fluorophores may blink many times, only once, or not at all (47). This phenomenon makes it difficult to distinguish nearby molecules of the same type. We employed an aggressive grouping algorithm to address this issue, but the tradeoff is that true separate molecules may be missed. We labeled our own secondary antibodies to control the dye:antibody ratio at ~1:1 to limit multiple blinking, but the labeling creates a distribution and some antibodies will still have multiple fluorophores. Antibodies engineered to have consistent labeling stoichiometry would be an improvement. A tradeoff to limiting the dye:antibody ratio is that many of the secondary antibodies will have no fluorophore, reducing labeling efficiency. SMLM techniques generally have shown a labeling efficiency of at most 60% (26). Improvements to labeling efficiency will enable rare binding interactions to be more readily detected by our method.

A third challenge for this algorithm comes from drift. Autocorrelation drift correction is standard, but it is optimal for defined structures that can be aligned from repeated blinks that occur throughout the acquisition. Singular soluble proteins, which blink only within a small window, pose a challenge for this correction method, and too few blinks overall can prevent the automatic correction from working despite apparent drift by eye. Further, this correction method cannot remove high-frequency variation in position through vibration within the microscope. Fluorescent beads may be used as a fiducial marker at the coverglass surface, but a solution is needed that would work throughout the cell. A sparsely labeled ubiquitous cellular structure, like tubulin, could serve this purpose with an appropriately engineered label. Eliminating drift effects would allow this algorithm to use smaller radii, increasing its sensitivity.

Despite these challenges, our technique provides a significant advance over existing methods in providing single-molecule-resolution information on protein and RNA associations inside the native cellular context. Desire for an SMLM technique using a nearest neighbor approach has been discussed recently (26). Proximity ligation assays are also a comparable technique. In that technique, adjacent oligonucleotide-labeled antibodies are in situ amplified by rolling circle amplification and labeled with a fluorescent oligonucleotide probe (48). The strength of a PLA is that associated molecules are specifically identified in a cell, to the exclusion of all non-associated members, allowing a clear count of interactions. However, information on the total number of molecules is lost, and the resolution is limited. In contrast, SMLM enables the associations to be placed in the context of their whole cellular complement and can distinguish interactions between molecules located closer than the diffraction limit. Conceivably, one could develop a correlative SMLM-PLA approach to combine their strengths.

Previously, we found that MEG3 induces p53 stabilization and stimulates p53–dependent transcription activation (16). In this study, we demonstrated that MEG3 lncRNA interacts with p53 inside the cell and can be detected with a novel analytical method using dSTORM. We also demonstrated that the p53–Mdm2 interaction may not be significantly disrupted by MEG3 in cells. Taken together, these data suggest that an alternative mechanism leads to p53 activation. Finally, we believe our association analysis provides a powerful new tool to assess macromolecular interactions in a native cellular context, with future extensions to 3-dimensional data and multi-protein complexes.

## Experimental procedures

### Cell lines, media, and growth conditions

The U2OS osteosarcoma cells (ATCC HTB-96) were maintained in Dulbecco’s modified Eagle’s medium (DMEM) (Gibco 11995065) supplemented with 10% heat-inactivated FBS (Gibco A3060502), glutamine (2 mM), penicillin (100 U/mL), and streptomycin (0.1 mg/mL) (Gibco 10378016) at 37 °C and 10% CO_2_. Doxycycline (Dox; 1 μg/mL) was added to media for at least 20 h to induce expression of the transfected tetracycline-inducible *MEG3.* Nutlin-3a (Nut; 10 μM) was added to media for at least 24 h to inhibit Mdm2-mediated degradation of p53. For microscopy, 3-5×10^4^ cells were seeded into each well of a chambered 8-well 1.5H coverglass (Ibidi 80827) and allowed to adhere overnight prior to further manipulation. Cells were tested for mycoplasma contamination every three months. U2OS-MEG3 cells were regularly authenticated by qRT-PCR and/or FISH for induction of MEG3 by doxycycline.

### Plasmid construction and transfection

A modified Tet-On expression system was used to express MEG3, consisting of pBiTetO-MEG3-GFPLoxP and pCMV-rtTA3-IRESpuro. pBiTetO was constructed by replacing the CMV promoter in expression vector pCI with a tetracycline responsive bi-directional promoter, BiTetO, which was synthesized to contain 7 modified TetO elements flanked by two minimal CMV promoter sequences based on pTet-T2 sequences (GenScript) (49). To facilitate selection of clones, a GFP cDNA with the coding region flanked by two LoxP sites was cloned into pBiTetO to generate pBiTetO-GFPLoxP. The *MEG3* cDNA in pCI-MEG3 (16) was modified by replacing AATAAA and its downstream poly(A) tail with a genomic DNA fragment containing the *MEG3* gene polyadenylation signal. The resultant *MEG3* cDNA was then cloned into pBiTetO-GFPLoxP to generate pBiTetO-MEG3-GFPLoxP. To construct pCMV-rtTA3-IRESpuro, a modified tetracycline responsive transactivator (rtTA3) was synthesized with changes in three amino acids including F67S, F86Y, and A209T (16,50) and inserted into pIRESpuro3 (Clontech Laboratories). Plasmids were verified by sequencing.

For stable transfection, U2OS cells were seeded into 6-well cell culture plates and transfected with pBiTetO-MEG3-GFPLoxP and pCMV-rtTA3-IRESpuro at a ratio of 3 to 1 using Mirus TransIT-LT1 according to the manufacturer’s instructions. Forty-eight hours after transfection, cells were reseeded in P100 dishes with limited dilution. Approximately ten days after treatment with puromycin (2 μg/mL), drug resistant colonies were isolated using cloning rings. Cells from individual clones were treated with or without doxycycline (1 μg/mL) for 24 h. GFP expression was evaluated under a fluorescence microscope. Cells expressing GFP in Dox-treated wells were further examined for MEG3 expression by qRT-PCR. Two sets of primers were used to detect MEG3. The first set detected a fragment near the 5’ end of the MEG3 cDNA: 5’-ATTAAGCCCTGACCTTTGCTATGC-3’ (forward) and 5’-ATAAGGGTGATGACAGAGTCAGTCG-3’ (reverse); the second set detected the 3’ end of the MEG3: 5’-CTTCAGTGTCTGCATGTGGGAAG-3 ‘ (forward) and 5’-TGCTTTGGAACCGCATCACAG-3’ (reverse). The GAPDH gene was used as an internal reference. The primers for detection of GAPDH were: 5’-GATGACATCAAGAAGGTGGTGAAGC-3’ (forward) and 5’-CGTTGTCATA-CCAGGAAATGAGCTTG-3’ (reverse). Cell clones with suitable MEG3 induction were treated with adenoviruses expressing Cre (Ad-Cre) to remove the floxed GFP. Up to three rounds of virus treatments were needed to completely remove GFP. The removal of GFP was confirmed by qRT-PCR with primer set: 5’-CCACAACGTCTATATCATGGCCG-3’ (forward) and 5’-GTGCTCAGGTAGTGGTTGTCG-3’ (reverse). A total of four clones containing inducible MEG3 were finally obtained and designated as U2OS-MEG3.

### Direct stochastic optical reconstruction microscopy (dSTORM)

#### Fixation

Cells were grown to between 30-90% confluence in chambered coverglass. Cells were rinsed with prewarmed Dulbecco’s phosphate-buffered saline with calcium and magnesium (DPBS; Corning) twice using near-simultaneous aspiration and injection of liquid to avoid dehydration. Prewarmed fixation buffer (4% (v/v) paraformaldehyde (Electron Microscopy Sciences), 0.1% (v/v) glutaraldehyde (Electron Microscopy Sciences)) was added and incubated in the dark for 15 min. Fixed cells were rinsed with DPBS. Remaining fixative was quenched with 1% (w/v) sodium borohydride (Sigma-Aldrich) for 7 min. (0.1% is typical, but we have observed better suppression of autofluorescence at 1%.) Cells were further quenched and washed with 50 mM glycine (Bio-Rad) in DPBS (DPBS-G) 3 times for 10 min each. Fixed cells were stored for up to a week in DPBS at 4 °C.

#### Immunofluorescence

Cells were permeabilized with 0.2% Triton X-100 (*t*-octylphenoxypolyethoxyethanol, Sigma-Aldrich) in DPBS for 10 min and rinsed with DPBS. Cells were blocked with 5% normal donkey serum (EMD Millipore)/0.02% (v/v) Triton X-100 in DPBS for 4 h at room temperature or overnight at 4 °C. Primary antibodies (rabbit anti-p53 [7F5] (Cell Signaling 2527S, Lot 8), mouse anti-Mdm2 [2A10] (Abcam ab16895, Lot GR324625-5)) were applied at 1:1000 and 1:200 dilutions, respectively, in blocking buffer and incubated overnight at 4 °C. Cells were washed with DPBS 6 times for 5 min each. Secondary antibodies (donkey anti-rabbit IgG (Jackson ImmunoResearch) and donkey antimouse IgG (Jackson ImmunoResearch)) were labeled as previously described with ATTO 488 (ThermoFisher Scientific) or Alexa Fluor 647 (ThermoFisher Scientific) for a dye ratio of ~1:1 (51). Secondary antibodies were added at 3 μg/mL each in blocking buffer and incubated for 2 h at room temperature in the dark. All subsequent steps were performed in the dark. Cells were washed with DPBS 6 times for 5 min each. Antibody stacks were crosslinked by 4% (v/v) paraformaldehyde in DPBS for 15 min. Remaining fixative was quenched and washed with DPBS-G twice for 5 min each, followed by DPBS twice for 5 min each. Stained cells were stored at 4 °C for up to 2 weeks before imaging.

#### Combined immunofluorescence and fluorescence in situ hybridization (FISH)

All buffers are RNase-free. Cells were permeabilized with 0.2% Triton X-100 in RNase-free PBS (Corning) for 10 min and rinsed with PBS. No blocking was performed to avoid introducing RNase activity. Primary antibody (rabbit anti-p53, see above) were applied at 1:1000 or 1:200 dilutions, respectively, in PBS and incubated overnight at 4 °C. Cells were washed with PBS 6 times for 5 min each. FISH was performed using buffers and ~20-mer tiled probe sets from Stellaris, according to manufacturer’s protocol. In brief, cells were washed with Wash Buffer A 2 times for 3 min. MEG3-Quasar 670 (Stellaris, custom order) or GAPDH-Quasar 570 (Stellaris SMF-2026-1) probe mixture was mixed 1:1000 in Hybridization Buffer and 100 μL was added per well. Steps from this point forward were conducted in the dark. The chambered coverglass was placed in a pre-warmed humidified chamber (large culture dish with damp paper towels) and incubated at 37 °C for 16 h. Cells were washed 2 times for 15 min each with warm Wash Buffer A in the humidified chamber. Secondary antibodies (donkey anti-rabbit conjugated with ATTO 488, see above) were added at 3 μg/mL each in Wash Buffer A and incubated for 1 h at 37 °C in the humidified chamber. Cells were washed 2 times for 2 min each with Wash Buffer B, then 2 times for 5 min each with PBS.

#### Imaging

Imaging buffer containing 10 mM cysteamine (2-mercaptoethylamine, MEA; Sigma-Aldrich), 3 U/mL pyranose oxidase from *Coriolus* sp. (Sigma P4234), and 90 U/mL catalase was freshly prepared in STORM buffer. Cysteamine stock solution was previously titrated to pH 8 and aliquots frozen. Precipitate in pyranose oxidase/catalase 100x enzyme stock solution was cleared by centrifugation at over 14,000×*g* prior to use. STORM buffer was composed of 10% (w/v) glucose, 10 mM sodium chloride, and 50 mM Tris hydrochloride (pH 8.0). We found the pyranose oxidase buffer (first described in (52)) to be superior to the standard glucose oxidase buffer. This buffer allowed longer imaging times due to minimal pH change, and the enzyme stock lasted several months at 4 °C with no observable decline in imaging quality. 10 mM cysteamine was selected for superior imaging characteristics with different dyes (53). PBS was replaced with the imaging buffer and the slide was mounted on the stage with type F immersion oil (refractive index = 1.515) on a Nikon Ti2 Eclipse inverted microscope. The microscope was equipped with a 100× 1.49 NA APO-TIRF objective with automatic correction collar and a Nikon NSTORM system including 405 nm (20 mW), 488 nm (70 mW), 561 nm (70 mW), and 647 nm (125 mW) lasers, a quadband excitation-emission filter, and a Hamamatsu ORCA Flash4.0 V2 S-CMOS camera. Nikon Elements 5.02 was used for image acquisition. A 10×10 tiled (with 15% overlap) widefield fluorescence image (~790×790 μm^2^) was obtained with 1 s exposure using GFPHQ, TexasRedHYQ, or Cy5HYQ filter cubes, from which random individual cells were selected for dSTORM imaging. At least 11000, 256×256 pixel (160 nm/pixel) frames were collected with 10 ms exposure time at 100% laser power with lasers in highly inclined and laminated optical sheet (HILO) configuration (54). Each channel was collected sequentially from longest wavelength to shortest.

### Data analysis

Localizations were identified from STORM image stacks using Nikon Elements 5.02 (NSTORM 4.0), with a peak height threshold of 250. Localization lists were exported as tab-delimited text files.

Localization data was processed with custom code written in the freely available Julia scientific computing language (v1.6) (55). Localizations identified in the first 100 frames, while fluorophores are being placed into the “off” state, were excluded. Localizations identified in the last 10 frames were also excluded due to artifacts caused by the change in optical configuration. For each image, a grouping algorithm (SI Algorithm 1; Figure S9) was applied to each channel to combine repeated blinking from single fluorophores and localizations that may be associated (e.g., another fluorophore on same secondary antibody, another secondary antibody on the same primary antibody, another primary antibody on a multimer). The first stage of the grouping algorithm iteratively identified local density maxima by searching a 34.2 nm radius and within a temporal window of 500 frames (5.0 s) of each localization for the localizations with the most neighbors, combining those localizations within the radius of the maxima, and repeating until all localizations were assigned to a group. The 34.2 nm radius limit was derived from a simulation of the possible orientation and positions of fluorophores in an antibody stack, to account for possible motion of the antibody stack and multiple fluorophores on the stack. The temporal window was applied to account for longer-scale on/off cycles of the fluorophores, as first described for PALM data (47), and was chosen semi-empirically by testing a range of values and selecting the smallest value that merged the most localizations (i.e., where the slope starts to decrease before the plateau) (Figure 10A) and where the merge results appeared suitable (e.g., few temporally separated clusters of localizations were merged).

**Figure 10.**
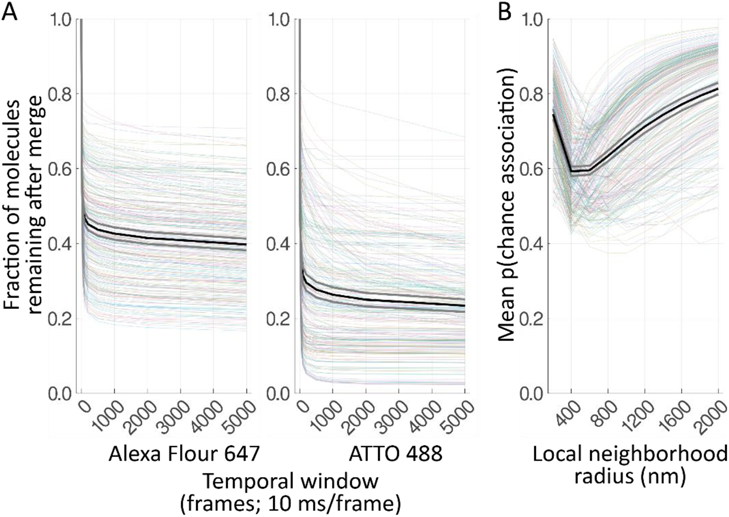
Analysis of algorithm parameters. All cells were repeatedly analyzed with the algorithm during development to characterize the effect of parameter choice on outcome. (A) Fraction of molecules remaining after grouping, normalized to the number of molecules with window size 1, as a function of temporal window size (frames, at 10 ms/frame), for Alexa Fluor 647 and ATTO 488. (B) Mean probability of chance association (p(chance association)) as a function of local neighborhood radius (nm). Each faded colored line is from a single cell. Black line shows the mean, gray lines indicate ± 95% confidence interval.

In the second stage, grouped localizations were merged if they were found within 200 nm of each other by a similar local density maxima search algorithm to further reduce redundancy from autocorrelated localizations. The products of this grouping algorithm were termed “molecules.” The position of the resulting molecule was the mean of its component localizations’ positions, and its linear localization accuracy was the mean of the accuracy for its component localizations divided by the square root of the number of component localizations.

The molecules were paired between channels by an exclusive cross-nearest neighbor algorithm (i.e., closest pair found and then removed, next closest pair found and then removed...; SI Algorithm 3) to obtain a distance distribution between the two sets of molecules. Two analytical approaches were applied, simple and sophisticated. First, the median paired distance was calculated for each cell. Second, a novel approach was developed to control for local density, based on a similar approach applied to single-molecule conventional fluorescence microscopy (29) but with significant modifications to address the different characteristics of SMLM data. Random permutations (10,000) of the molecules in the local (800 nm radius) neighborhood around each potential binding pair were generated and the closest pairwise distance in each permutation was calculated to create a Monte Carlo estimation of the distribution of distances due to local density (SI Algorithm 4). The local neighborhood radius of 800 nm was semi-empirically chosen based on testing multiple window sizes with the algorithm and choosing the value that provided a balance of sensitivity (smaller value) and robustness (less inter-sample change as parameter changes) (Figure 10B). The fraction of permutations with a closest distance less than the observed distance was the percentile rank score, indicating the probability of chance association given the local density of both molecule types. Finally, the fraction of pairs within the maximum binding distance (200 nm) and with a probability of chance association of less than 0.1 was calculated for each cell. The maximum binding distance was chosen based on knowledge of the size of the target molecules (up to 20 nm across) and the size of the antibody stacks (up to 70 nm), with allowance for error.

Data are shown as mean ± standard deviation. Data were tested for significance by ANOVA with replicates as a repeated measure, as we observed correlation of values by replicate, with α = 0.05, using the SimpleANOVA.jl (v0.8.0) Julia package created by the authors. Generalized ω^2^ effect size was interpreted as small (~0.01), medium (~0.06) and large (~0.14). Data was checked for extreme outliers, heteroscedasticity, and normality of residuals, and were determined to be reasonable. Single outlying datapoints with a z-score ≳ 2.25 in each sample were Windsorized. Plots were generated with StatsPlots.jl (v0.14.0) and assembled with Adobe Illustrator (24.0).

## Code availability

All the code generated specifically for this manuscript is written in the Julia language and available in the repository at doi: 10.5281/zenodo.4542449. Supporting packages can be obtained within Julia from its public package registry.

## Data availability

Raw STORM data files are stored on a local server. STORM localization list text files are available in the repository at doi: 10.5281/zenodo.4542454.

## Acknowledgements

We thank Dr. Angie Schmider and Nikon for training on the STORM system; Dr. Hongjae Sunwoo for advice on combining FISH and STORM; Dr. Carolina Eliscovich for help understanding her Monte Carlo-based method; and Dr. Hang Lee of Harvard Catalyst for statistical advice.

## Author contributions

N. C. B., Y. Z., A. K. and R. J. S. conceptualization; N. C. B. and Y. Z. data curation; N. C. B. formal analysis; N. C. B., A. K. and R. J. S. funding acquisition; N. C. B., A. Y., X. W. and Y. Z. investigation; N. C. B. and Y. Z. methodology; Y. Z., A. K. and R. J. S. project administration; A. Y., X. W. and Y. Z. resources; N. C. B. software; Y. Z., A. K. and R. J. S. supervision; N. C. B. and Y. Z. validation; N. C. B. visualization; N. C. B., Y. Z. and R. J. S. writing – original draft; N. C. B., Y. Z. and R. J. S. writing – review & editing.

## Funding and additional information

RJS: R01CA193520, R01DK062472, S10RR027931; NCB: T32DK007540; AK and YZ: R01CA193520; and the Jarislowsky Foundation. Stochastic optical reconstruction microscopy experiments were conducted at the MGH Molecular Imaging Core. The content is solely the responsibility of the authors and does not necessarily represent the official views of the National Institutes of Health.

