## Supplemental Information for "A cross–nearest neighbor/Monte Carlo algorithm for single-molecule localization microscopy defines interactions between p53, Mdm2, and MEG3"

### Algorithms

*Algorithm 1 – Group localizations into molecules*

1. Find the neighbors of all localizations within a radius of 34.2 nm and a temporal window of 500 frames (0.5 s) (Figure S9A).
2. Identify local density maxima for each localization (Algorithm 2) (Figure S9B).
3. Remove local density maxima from the localizations list and filter the dictionary mapping each localization to its maximum to exclude local maxima.
4. Create “molecules” from the local maxima.
5. Iteratively join the local maxima to all associated localizations within the 34.2 nm radius, plus the linear localization accuracy for each (Figure S9C and S9D).
6. Remove all joined localizations and go to 1 if any unjoined localizations remain (Figure S9E).
7. Merge molecules by going to 1 using the molecules as the source without the temporal window and using a 200 nm radius (Figure S9F and S9G). Repeat until no more molecules can be merged.

*Algorithm 2 – Identify local density maxima*

1. Take a localization
2. Pick the neighbor (inclusive of this localization) with the most neighbors of its own
3. If the picked neighbor is not this localization, record the localization as visited and go to 2.
4. If the localization with the most neighbors is this one, add it to the list of local maxima, record in a dictionary that all previous localizations in this loop are associated with this local density maximum, and go to 1 if any unvisited localizations remain.
5. Return a list of local density maxima and a dictionary that associates all localizations with their local density maximum. All input localizations will either be a local density maximum or associated with one.

*Algorithm 3 – Exclusive cross-nearest neighbors*

1. Find the nearest neighbors between each set of molecules (NearestNeighbors.jl).
2. Extract the closest unique pairs and the distance between them.
3. Go to 1 if there are any unpaired molecules in the smaller of the two sets.
4. Return the list of pairs and distances.

*Algorithm 4 – Monte Carlo affinity*

1. For each molecule pair, count the neighbors of each type within 800 nm of the pair’s centroid.
2. For each unique count of each molecule type, generate 10,000 permutations of their positions within the 800 nm radius and calculate the distance of the closest observed pair in each, stored in a dictionary.
3. For each molecule pair, calculate the percentile rank score as the fraction of permutations generated in 2 in which a smaller distance was obtained than the distance observed.
4. Return the percentile rank score for each molecule pair.

**Figures**

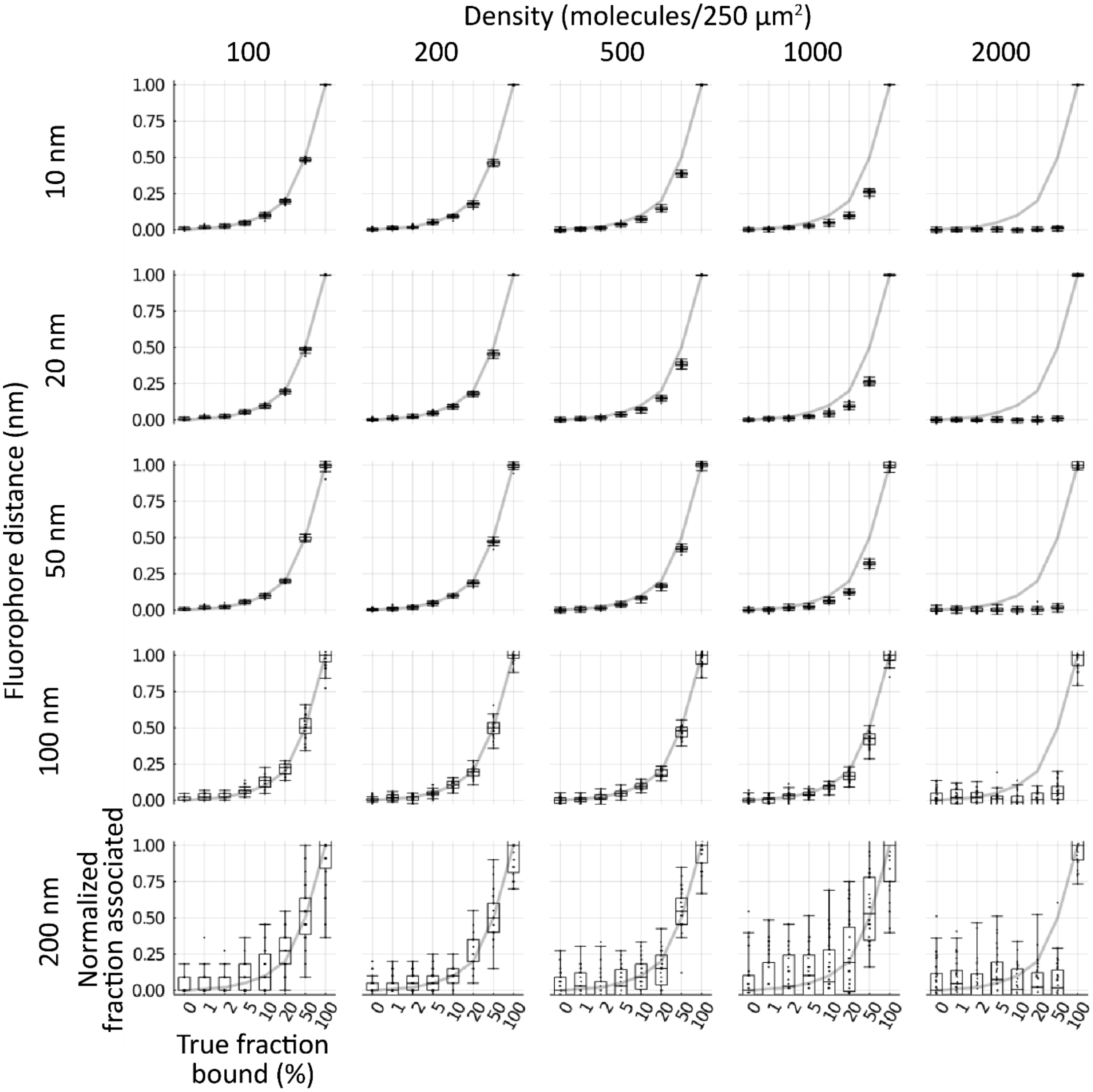


**Figure S1. Fraction associated of simulated data normalized to 0% and 100%.** Simulated cells of 100, 200, 500, 1000, 2000 molecules of each type (columns left to right) within a 250 µm$^2$ circle with 10, 20, 50, 100, 200 nm separation between pairs (rows top to bottom) for cells with 0, 1, 2, 5, 10, 20, 50, or 100% binding were generated (n = 30 each condition). Cells were run through the cross–nearest neighbor/Monte Carlo algorithm and then normalized to the mean of the 0% and 100% samples in each condition. Boxes indicate median +/- upper and lower quartile; whiskers indicate the range excluding outliers. The gray line indicates the expected values at each point.

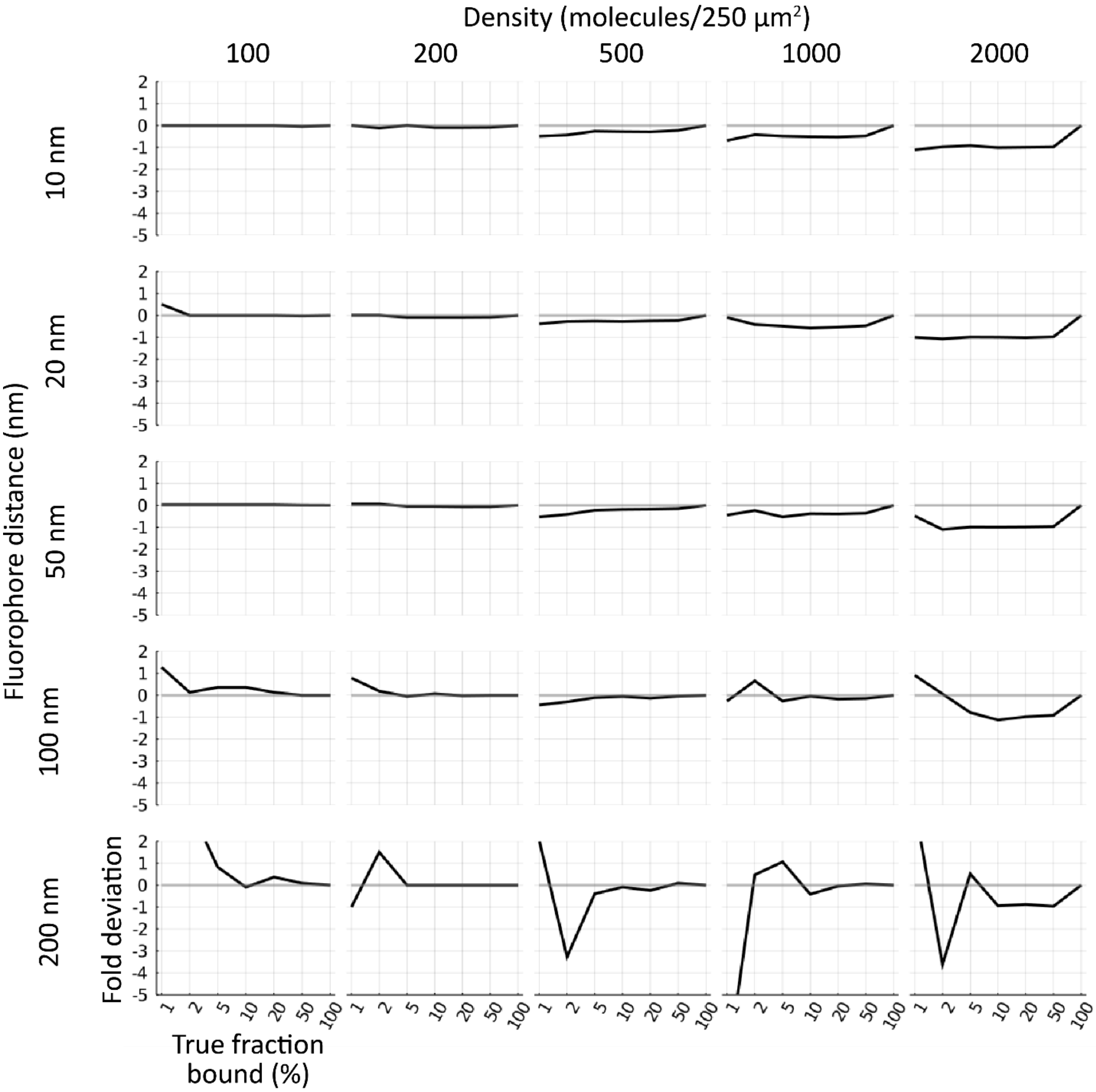


**Figure S2. The measured value underestimates the true value substantially at high density and large distances between fluorophores.** Simulated cells of 100, 200, 500, 1000, 2000 molecules of each type (columns left to right) within a 250 μm$^2$ circle with 10, 20, 50, 100, 200 nm separation between pairs (rows top to bottom) for cells with 0, 1, 2, 5, 10, 20, 50, or 100% binding were generated (n = 30 each condition). Cells were run through the cross–nearest neighbor/Monte Carlo algorithm and then normalized to the mean of the 0% and 100% samples in each condition. The line indicates the relative deviation from the true value (negative indicates an underestimate).

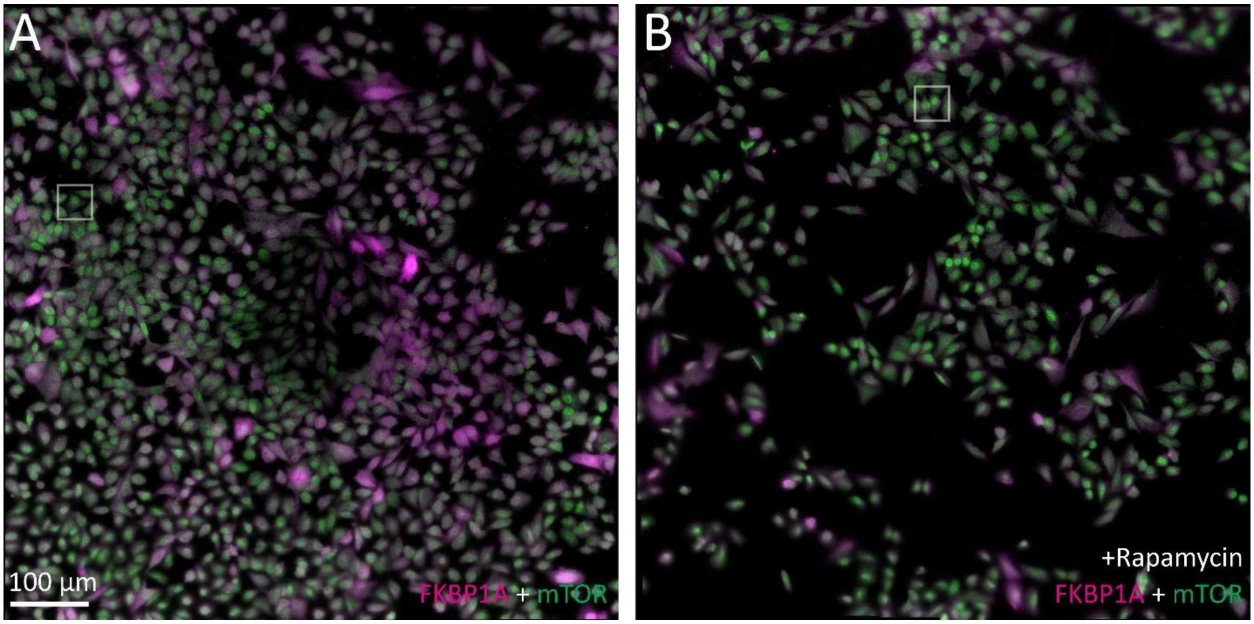


**Figure S3. Representative tiled micrographs of U2OS cells containing doxycycline-inducible MEG3 stained for FKBP1A and mTOR.** Cells were treated for 24 h with (B) or without (A) rapamycin. Cells were stained for FKBP1A with an Alexa Fluor 647 secondary antibody (magenta) and mTOR with an ATTO 488 secondary antibody (green). All images composed of 10x10 micrographs stitched together with 10% overlap and 1-second exposures. Scale bar is 100 µm. White box indicates the representative cells shown in the text.

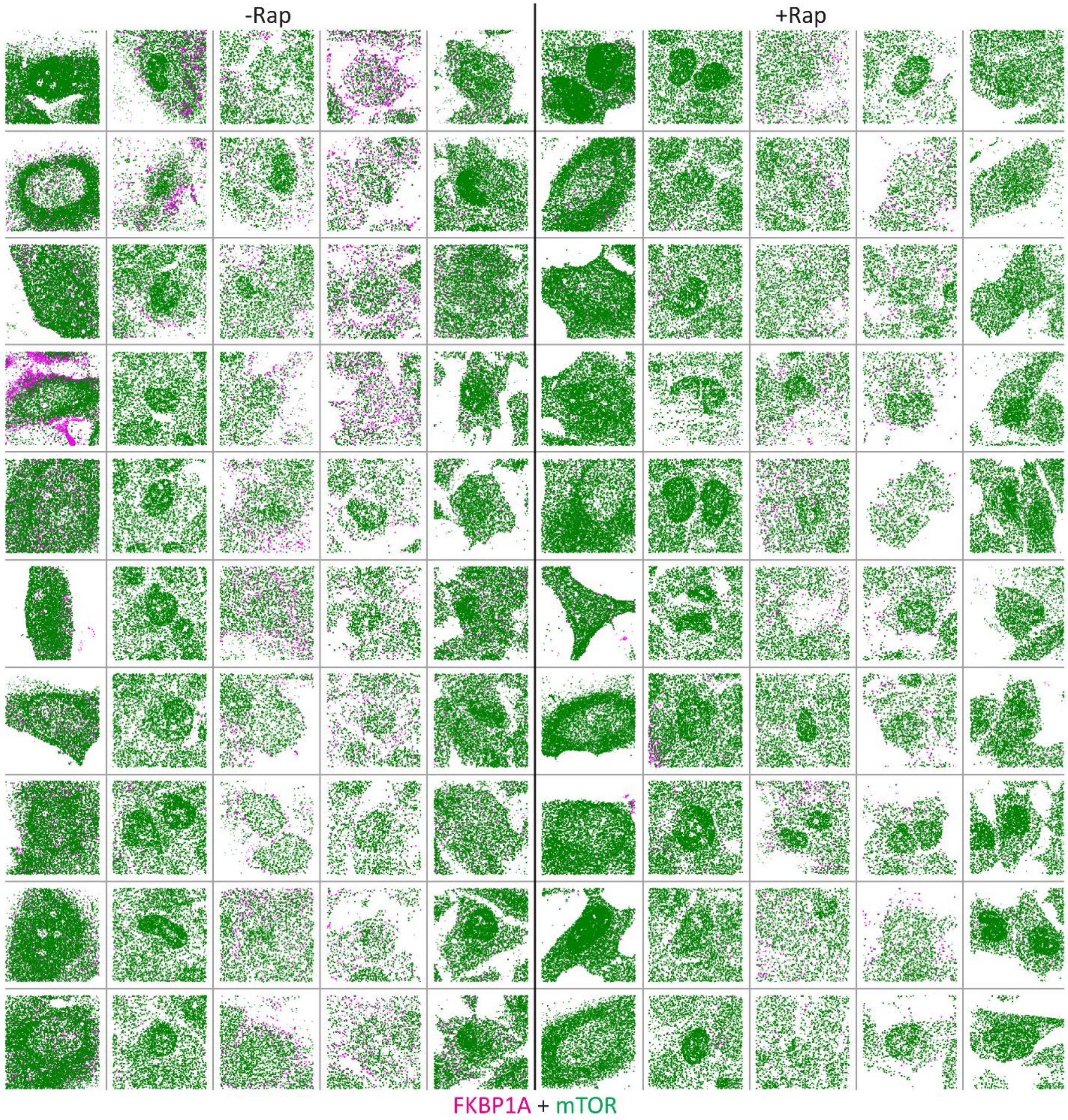


**Figure S4. Localization maps of cells used in analysis of FKBP1A and mTOR association.** Cells were treated for 24 h with (right) or without (left) rapamycin. Cells were stained for FKBP1A with an Alexa Fluor 647 secondary antibody (magenta) and mTOR with an ATTO 488 secondary antibody (green) and imaged by dSTORM. Each column is from one replicate. Each square is 40.960 µm across.

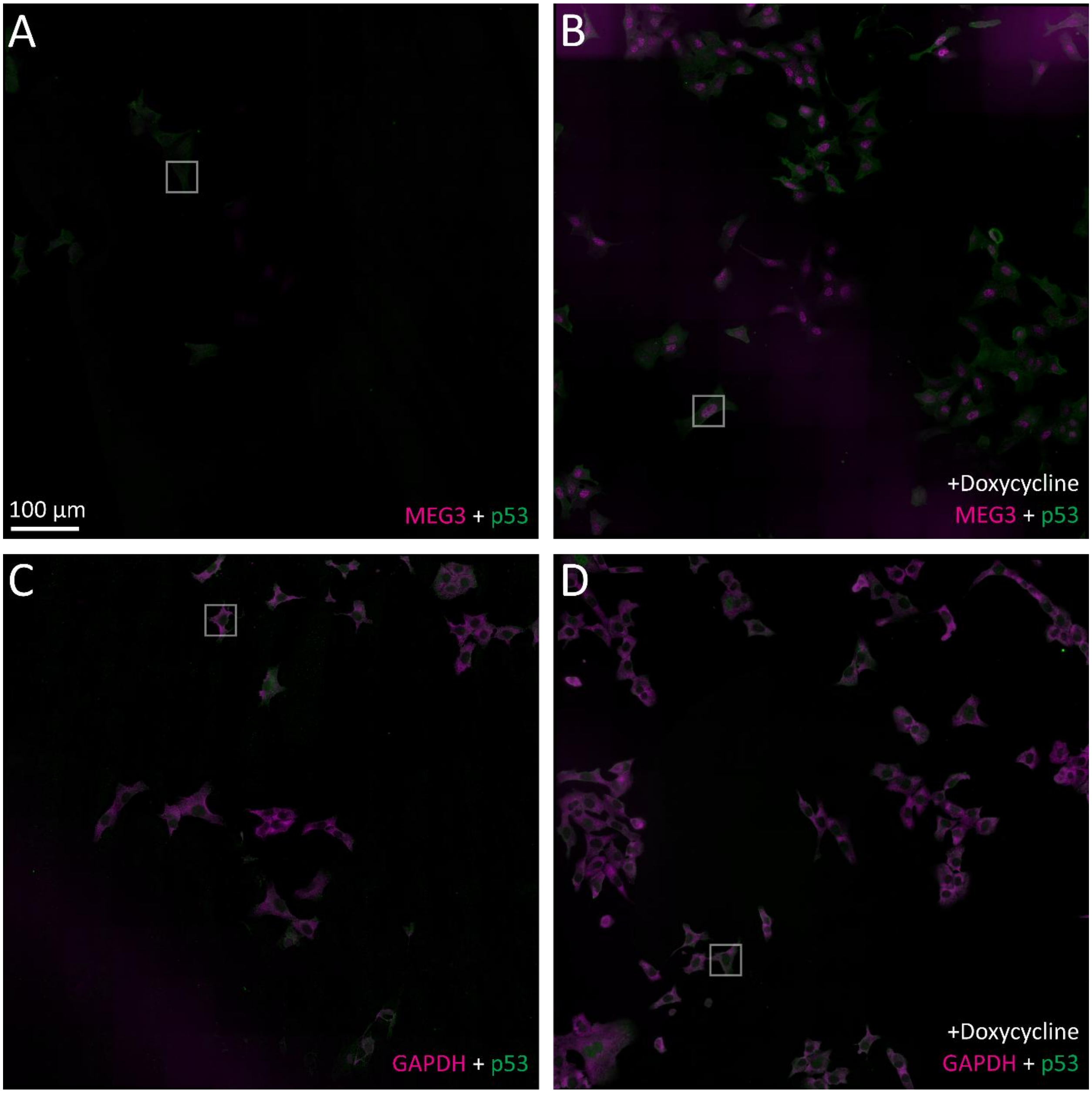


**Figure S5. Representative tiled micrographs of U2OS cells containing doxycycline-inducible MEG3 stained for p53, MEG3, and *GAPDH* mRNA.** Cells were induced for 20 h with (B, D) or without (A, C) doxycycline. Cells were stained for p53 with an ATTO 488 secondary antibody (green) and FISH probes for either MEG3 (Quasar 670, magenta; A, B) or *GAPDH* mRNA (Quasar 570, magenta; C, D). All images composed of 10x10 micrographs stitched together with 10% overlap and 1-second exposures. Scale bar is 100 µm. White box indicates the representative cells shown in the text.

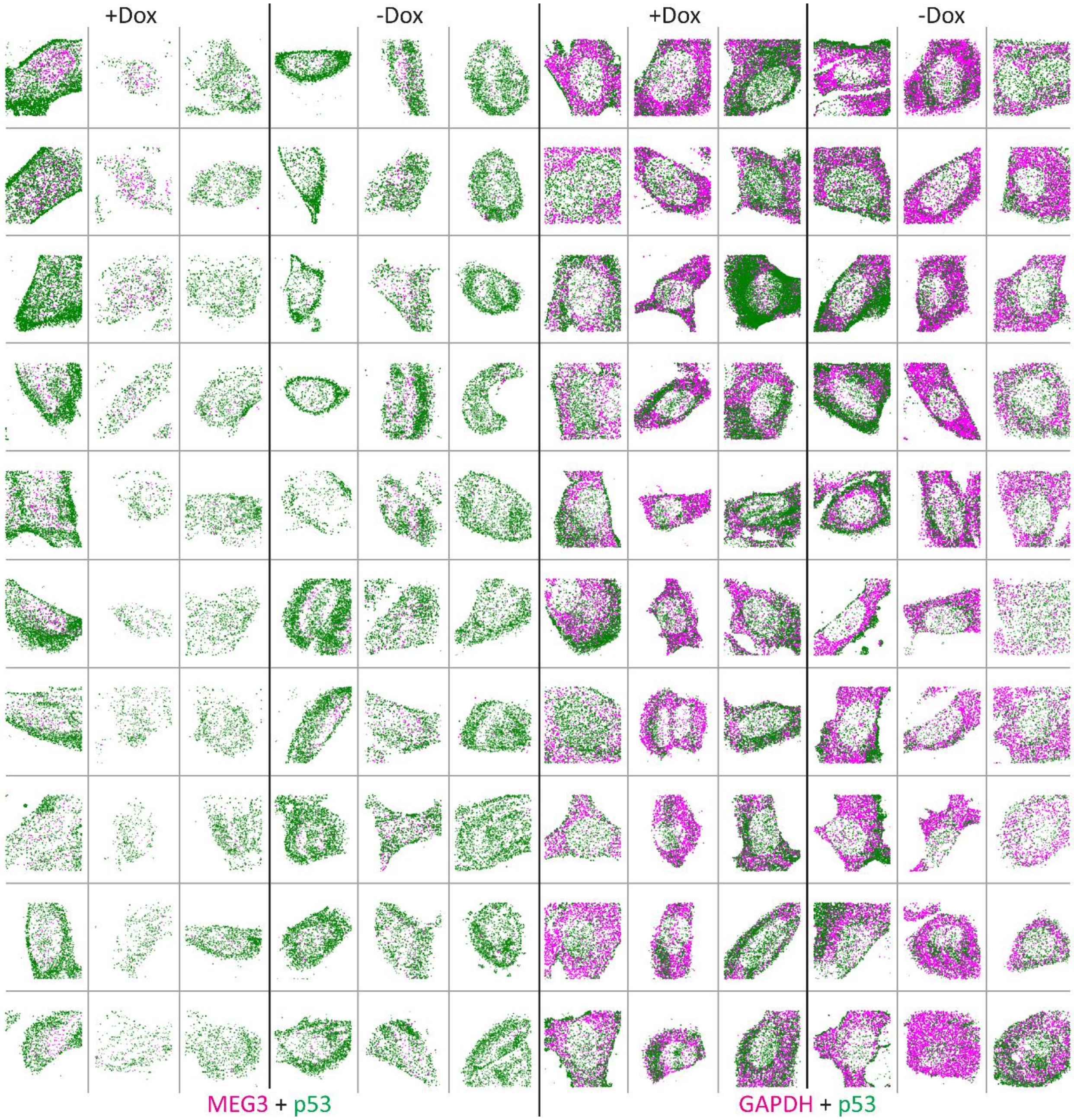


**Figure S6. Localization maps of cells used in analysis of MEG3, *GAPDH* mRNA and p53 association.** U2OS-MEG3 cells were induced for 24 h with (far left, middle right) or without (middle left, far right) doxycycline. Cells were stained for p53 with an ATTO 488 secondary antibody (green) and FISH probes for either MEG3 (Quasar 670, magenta) or *GAPDH* mRNA (Quasar 570, magenta) and imaged by dSTORM. Each column is from one replicate. Each square is 40.960 μm across.

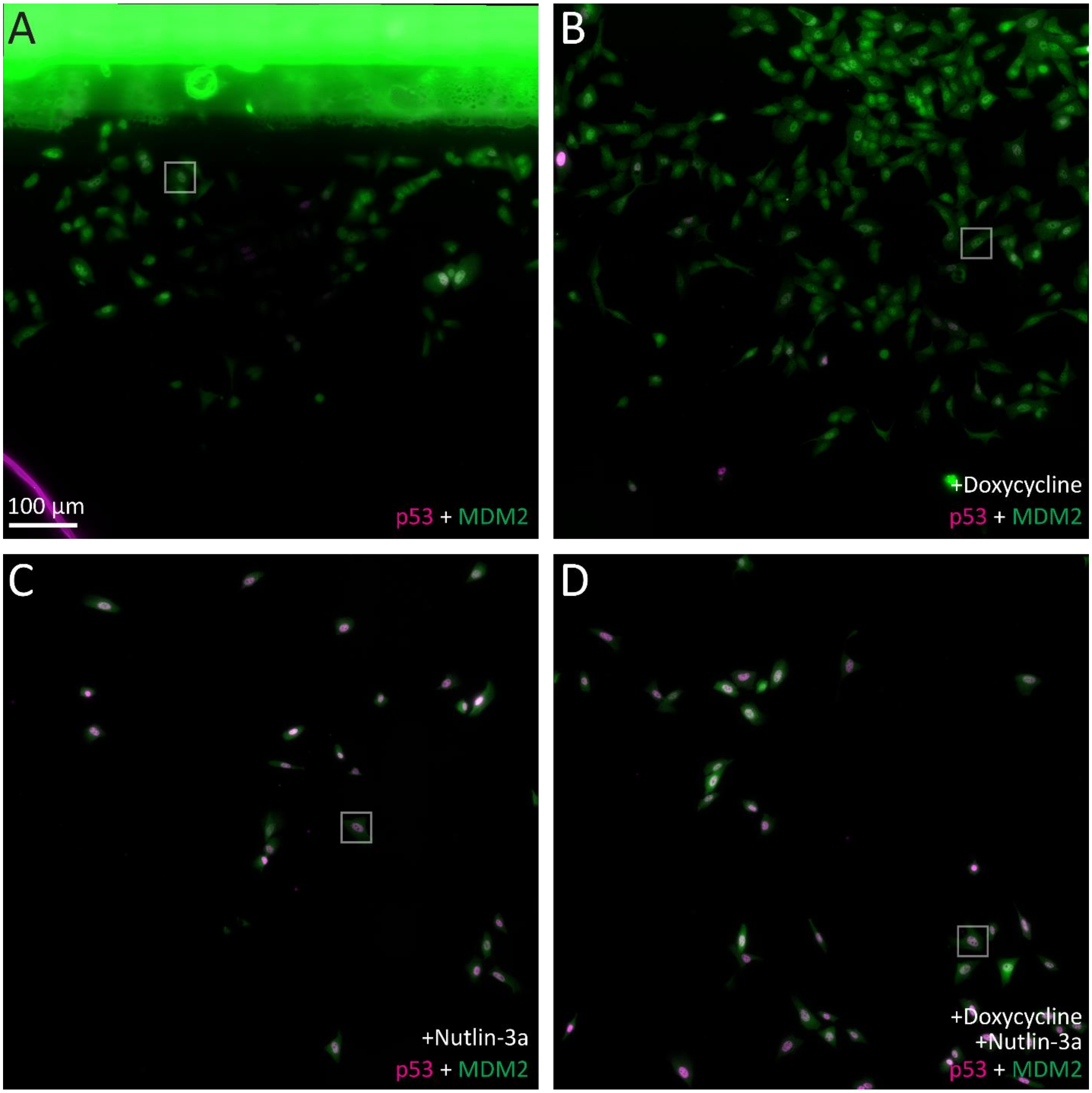


**Figure S7. Representative tiled micrographs of U2OS cells containing doxycycline-inducible MEG3 stained for p53 and Mdm2.** Cells were induced for 20 h with (B, D) or without (A, C) doxycycline or with (C, D) or without (A, B) nutlin-3a. Cells were stained for p53 with an Alexa Fluor 647 secondary antibody (magenta) and Mdm2 with an ATTO 488 secondary antibody (green). All images composed of 10x10 micrographs stitched together with 10% overlap and 1-second exposures. Scale bar is 100 µm. White box indicates the representative cells shown in the text.

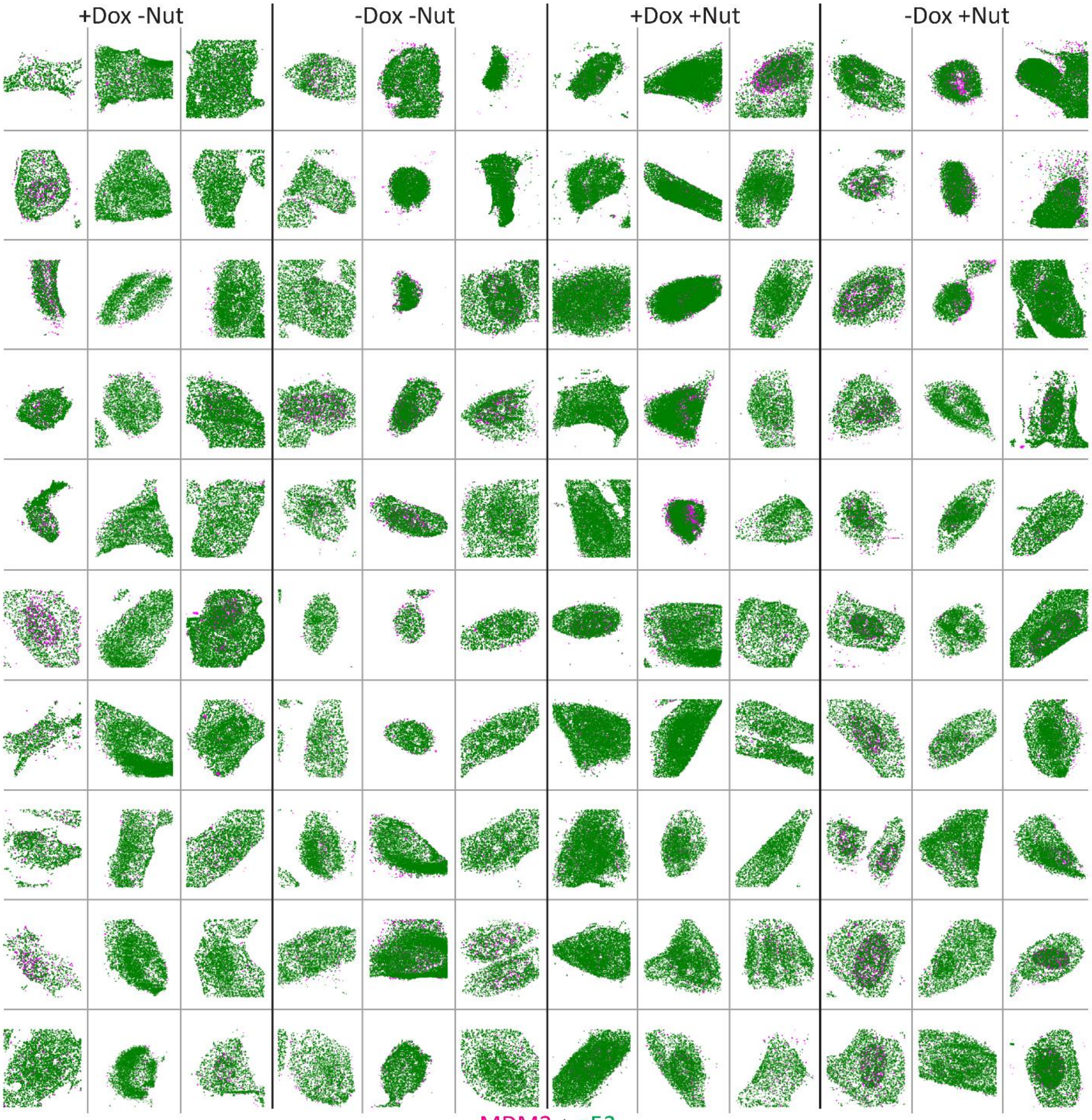


**Figure S8. Localization maps of cells used in analysis of Mdm2 and p53 association.** U2OS-MEG3 cells were induced for 24 h with (far left, middle right) or without (middle left, far right) doxycycline; and with (right) and without (left) nutlin-3a. Cells were stained for p53 with an Alexa Fluor 647 secondary antibody (magenta) and Mdm2 with an ATTO 488 secondary antibody (green) and imaged by dSTORM. Each column is from one replicate. Each square is 40.960 μm across.

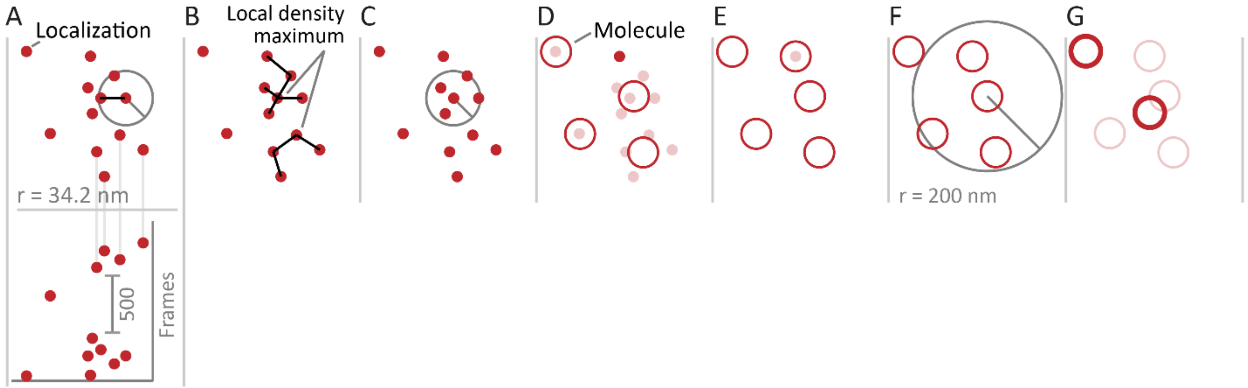


**Figure S9. Grouping algorithm.** (A) Localizations (red dots) are searched within a 34.2 nm radius and a 500-frames (0.5 s) temporal window for local density maxima. (B) Local density maxima are identified and localizations associated with them and within 34.2 nm (C) are joined into molecules with averaged positions (D). (E) The process repeats until all localizations are joined to molecules (red circles). (F) Nearby molecules are merged within 200 nm to reduce redundancy (G).
